# Stable Zn isotopes reveal the uptake and toxicity of zinc oxide engineered nanomaterials in *Phragmites australis*

**DOI:** 10.1101/2020.04.08.031179

**Authors:** C Caldelas, F Poitrasson, J Viers, JL Araus

**Affiliations:** Department of Evolutive Biology, Ecology, and Environmental Sciences. University of Barcelona. Av. Diagonal, 643, 08015, Barcelona, Spain; Géosciences Environnement Toulouse, UMR 5563 Centre National de la Recherche Scientifique - Université de Toulouse - Institut de Recherches pour le Développement, 14-16, avenue Edouard Belin, 31400, Toulouse, France; AGROTECNIO Center, University of Lleida, 25198 Lleida, Spain

## Abstract

The uptake, transport, and toxicity mechanisms of zinc oxide (ZnO) engineered nanomaterials (ZnO-ENMs) in aquatic plants remain obscure. We investigated ZnO-ENM uptake and phytotoxicity in *Phragmites australis* by combining Zn stable isotopes and microanalysis. Plants were exposed to four ZnO materials: micron-size ZnO, nanoparticles (NPs) of <100 nm or <50 nm, and nanowires of 50 nm diameter at concentrations of 0-1000 mg l^−1^. All ZnO materials reduced growth, chlorophyll content, photosynthetic efficiency, and transpiration and led to Zn precipitation outside the plasma membranes of root cells. Nanoparticles <50 nm released more Zn^2+^ and were more toxic, thus causing greater Zn precipitation and accumulation in the roots and reducing Zn isotopic fractionation during Zn uptake. However, fractionation by the shoots was similar for all treatments and was consistent with Zn^2+^ being the main form transported to the shoots. Stable Zn isotopes are useful to trace ZnO-ENM uptake and toxicity in plants.

**Environmental Significance Statement:** Our understanding of zinc oxide nanomaterials interaction with wetland plants is hampered by the lack of scientific consensus about their uptake and toxicity mechanisms in these species. This is a serious concern given the alarming global increase in the discharge of these nanomaterials into the environment and the key ecological roles of wetland plants. The Zn isotopic signature of plant tissue integrates all the Zn metabolic pathway throughout the plant’s life, giving insight about the form of Zn taken up, even if this later transforms into another Zn species. Thus, our findings clarify the exposure routes and the mechanisms of action of zinc oxide engineered nanomaterials in wetland plants while advancing the toolbox for plant physiology and environmental studies.

**Table of contents:** 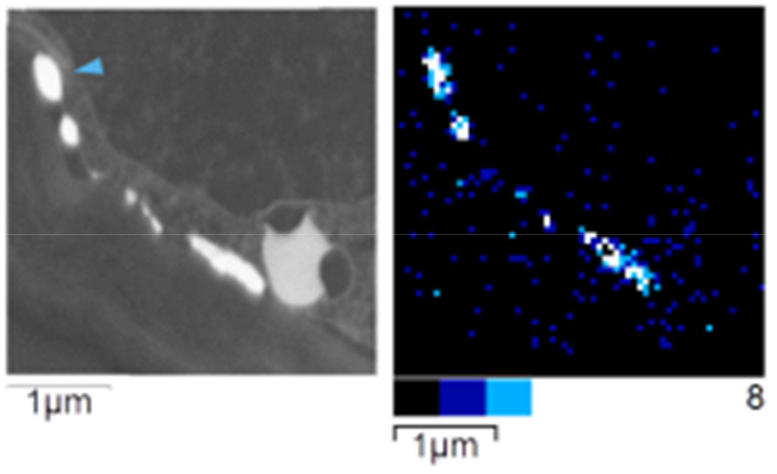

The Zn stable isotope composition of plants demonstrates that ZnO engineered nanomaterials dissolve before their uptake and accumulation by the roots (brightest inclusions in root cortex above).

## Introduction

An estimated 34,000 tonnes year^−1^ of ZnO-ENMs are emitted into the environment globally, of which 3,000 are directly discharged into water bodies^1^. Models predict that sediments will receive most of the ZnO-ENMs released into water bodies (up to 1 mg Kg^−1^ year^−1^)^2^, especially near large cities and related industry, in areas that favour deposition like wetlands and marshes. This may cause chronic toxicity in emergent wetland plants (helophytes), which play vital ecological roles^3^. Exposure to ZnO-ENMs decreases photosynthesis, antioxidant activity, and growth in aquatic plants, and increases Zn bioaccumulation and oxidative stress^4–10^. However, studies on helophytes are few and the uptake and toxicity of ZnO-ENMs, especially after chronic exposure, are poorly characterised in these plants.

There is an unresolved controversy about the capacity of entire ZnO-ENMs to enter roots. In several crops, ZnO-ENMs have been reported in the root epidermis, cortex, endodermis, stele, lateral roots, and on the root surface^11–16^. At the cellular level, ZnO-ENMs have accumulated in the intercellular spaces, along the plasma membrane, and in the cytoplasm, vacuoles and nuclei^11,14^. It has been suggested that ZnO-ENMs in the root apoplast can enter the symplast by endocytosis^11^ and pass to the xylem directly from the apoplast, through lateral roots with immature Casparian bands^6,14^. However, few studies have confirmed either the elemental composition of the candidate ENMs or the Zn speciation in roots. Wang *et al* found 65% ZnO and 32% Zn-histidine in the roots of hydroponically grown cowpea exposed to ZnO-ENMs^12^. Nonetheless, Da Cruz *et al* demonstrated that ZnO was only found inside roots when they had been previously damaged^13^. Accordingly, ZnO-ENMs normally do not reach the shoots unless roots are damaged^12–15^. Intact roots generally contain no ZnO but do contain other Zn species like Zn-phosphates, Zn-citrate, Zn-malate, Zn-histidine, or Zn-nitrate^13,14,16^. In the aquatic plant *Schoenoplectus tabernaemontani*, ZnO-ENMs were reportedly found around plastids in the roots, arranged like beads on a string^6^. However, this observation could be better explained as endoplasmic reticulum wrapped around plastids^17^.

The present research aims to give a comprehensive insight on ZnO-ENM uptake and chronic toxicity in *P. australis* at the levels estimated by models in current sediments, with an ample margin for a future increase in concentrations. This is crucial because emissions are expected to continue growing. To do so, we determined whether entire ZnO-ENMs enter *P. australis* roots by combining transmission electron microscopy (TEM), X-ray microanalysis, dark-field mapping, and Zn stable isotope analysis in different plant organs. Zinc stable isotopes are valuable to understand Zn uptake and toxicity in plants^18,19^. The Zn isotopic signature of plant tissue integrates all the processes causing Zn fractionation throughout a plant’s life and it is a proxy of Zn speciation and the activity of Zn membrane transporters^20^. Hence, Zn isotopes can be used to determine whether Zn was first taken up as either Zn^2+^ or ZnO-ENMs, even after they form other Zn species. Additionally, we unravelled the mechanisms of ZnO-ENM toxicity on the photosynthetic apparatus and fully characterised the physiological response of *P. australis* to ZnO-ENMs.

## Methods

### Nanomaterials and non-nano ZnO

Four ZnO materials were purchased from Sigma-Aldrich: non-nano ZnO (96479 Fluka, ACS reagent ≥99.9%); spherical particle nanopowder <100 nm in diameter (544906 Aldrich, ≈80%); <50 nm nanopowder particles of various shapes (677450 Aldrich, >97%); and nanowires 50 nm in diameter x 300 nm long (773980 Aldrich). These materials will be referred to as Bulk, NP100, NP50, and NW, respectively. To determine their chemical composition, 2 ml HNO_3_ (69.0-70.0% Instra reagent, 9598-34 Baker) was added to 50 mg aliquots and digested overnight at 90°C. The extracts were brought to 100 ml with 1% HNO_3_ and analysed for Al, Cd, Cr, Cu, Fe, Ni, Pb, and Zn content by inductively coupled plasma optical emission spectrometry (ICP-OES) using a Perkin Elmer Optima-8300 (Waltham, MA, USA). Three replicates were analysed for each ZnO material. To study particle size and shape, samples were observed on a field emission scanning electron microscope (FESEM) JEOL JSM 7100F at 20.0 kV, and secondary electron images were taken.

### Plant growth conditions

*Phragmites australis* (Cav.) Trin. Ex Steud plants were purchased from a local nursery (Tres Turons, Sabadell, Spain). Plant cultivation took place in the greenhouse of the Experimental Field Services of the University of Barcelona (UB). Roots were washed to remove the original substrate and shoots cut to induce new growth. Plants were initially grown in trays filled with modified half-strength Hoagland’s solution as detailed in^18^. The pH was adjusted to 6.5 and solution was replaced every three days. After 7 weeks, plants were rinsed in distilled water, selected within a small range of fresh weight (11.99 ± 0.23 g, mean ± standard error, n = 95) and height (27.54 ± 0.34 cm), and placed in individual 1-gallon (3.895 l) glass pots. These were filled with modified half-strength Hoagland’s solution with no added Zn according to^21^ (Supplementary Table 1), at pH 5.9. Each pot was wrapped in aluminium foil to limit algal growth. After 10 days, the ZnO materials described above were added at a concentration of 0, 0.1, 1, 10, 100, or 1000 mg l^−1^. Nanowires were only tested up to 10 mg l^−1^ due to their high cost. Four pots per treatment were distributed randomly on the greenhouse table, and the nutrient solution was replenished three times a week. Five pots without plants but containing the same solution were included. Plants were grown for 14 weeks, from November 2016 to February 2017. The mean temperature was 17.9±0.22°C, the relative humidity 51.0±1.06%, and the maximum PPFD (photosynthetic photon flux density) ~500 μmol m^−2^ s^−1^. The pH and electrical conductivity (EC) of the solutions were controlled weekly in a sub-sample of 10 pots (results not shown) and recorded for the whole experiment on weeks 8 and 12 (Supplementary Table 2).

### Elemental analyses

At the end of the experiment, 10 ml aliquots of the growth solutions were centrifuged for 10 minutes at 9065 relative centrifugal force in a Hettich 32R centrifuge (Tuttlingen, Germany) with a 1620A angle rotor. Supernatants were filtered using Whatman^®^ Grade GF/A glass microfibre filters (WHA1820042) and 100 μl HNO_3_ was added (Baker Instra-Analysed Reagent, 69.0-70.0%, 9598-34). The Zn and Al contents of the solutions were measured by ICP-OES.

Plants were thoroughly washed in distilled water, patted dry, and weighed. Roots and shoots were separated using a scalpel. Samples were oven-dried at 60°C for 48h and finely ground for 2 min in a Retzch MM400 mixer mill at a frequency of 20 s^−1^. Next, 0.1 g of the ground material was digested overnight at 90°C in a mixture of 2 ml HNO_3_ and 0.5 ml H_2_O_2_ (30% Suprapur, 1.07298.1000 Merck). Then 50 μl of HF (40% reagent grade, AC10511000 Scharlab) were added to each sample and digested for an additional 2h at 90°C. Digests were diluted in 30 ml MilliQ water (18.2 Ω) before analysis. Per every 30 samples, 3 blanks and 2 aliquots of the BCR-60 reference material (*Lagarosiphon major*) (Community Bureau of Reference, Brussels, Belgium) were digested using the same protocol. We obtained 303.1±5.3 (n=6) μg g^−1^ Zn, in line with the certified values (313.0±8.0). The Al, Ca, Cu, Fe, K, Mg, Mn, P, S, and Zn contents of the extracts were then determined by ICP-OES. To study Zn mass balance at the end of the experiment, Zn in mg was calculated for each reservoir in our culture system (Equation 1):

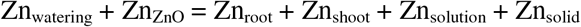

The Zn input contributed by the nutrient solution (Zn_watering_) was calculated by multiplying the [Zn] of the initial nutrient solution by the total volume of solution added and by the sum of all waterings (E). Zinc input from ZnO materials (Zn_ZnO_) was obtained from the [ZnO] of each treatment (0.1, 1, 10, 100, and 1000 mg l^−1^) multiplied by the Zn concentration of each material. Zinc extracted by roots (Zn_root_) and shoots (Zn_shoot_) was determined by multiplying [Zn] by dry weight (DW). The total Zn in the final solutions (Zn_solution_) was calculated by multiplying [Zn] in the final solutions by the volume of the container. Finally, the Zn in the solid phase of the nutrient media (Znsolid) was inferred from subtracting Zn_root_, Zn_shoot_, and Zn_solution_ from the sum of all Zn inputs. The same procedure was followed to calculate the Al mass balance.

### Zinc separation and isotopic analyses

Zinc isotope analyses were carried out in the facilities of the Géosciences Environnement, Toulouse, France. In the clean lab (ISO3), 100 mg of sample were weighed in Teflon beakers. Samples were digested in several steps: i) 24h at room temperature in 1.5ml HNO_3_ and 1 ml H2O2; ii) 24h on a hotplate at 80°C with the same mixture, then evaporated; iii) 24h at 80°C in a mixture of 1.2ml HF and 1.2 ml HNO_3_, then evaporated; and finally iv) 24h on a hotplate at 115°C in 20 drops of HCl and 10 drops of HNO_3_, and evaporated. The digests were then refluxed in 15 ml 10% HNO_3_ for Zn quantification in an ICP-MS Agilent 7500 (Santa Clara, USA). Three aliquots of BCR-60 were ground and processed in the same manner as the samples to quantify the Zn contribution of the stainless-steel jars and balls. We obtained 298.2±2.1 μg g^− 1^ Zn for the ground BCR-60 and 301.3±6.9 for the non-ground. These values are slightly lower than the certified 313.0±8.0 μg g^−1^ Zn, which is likely due to the cleanroom being a much cleaner environment than a regular chemistry laboratory. The sample residual was weighed and refluxed in 1 ml of 7N HCl + 0.001% H_2_O_2_ overnight at 40°C. HCl and HNO_3_ were double distilled, HF was 41-51% (Fluka A513-P500 suprapur) and H_2_O_2_ 30% (Merk 1.07298 suprapur). Plastic and Teflon material was acid-washed before use. Zinc separation was performed in Poly-Prep chromatography columns (BIO-RAD, 731-1550) containing 2 ml of anion exchange resin AGMP-1M (100-200 mesh, chloride form) (BIO-RAD, 1411841). The resin was first cleansed three times with 10 ml 18.2Ω water followed by 7 ml 0.5N HNO_3_, then conditioned with 6 ml 7N HCl + 0.001% H_2_O_2_ before loading the samples. Matrix elements and Cu were eluted with 30 ml 7N HCl + 0.001% H_2_O_2_. Iron (Fe) was eluted with 10 ml 2N HCl + 0.001% H_2_O_2_ and the resin was rinsed in 2 ml of 0.5N HNO_3_. Finally, Zn fractions were eluted on 8 ml of 0.5N HNO_3_ and evaporated. Three aliquots of the reference material BCR 281 (ryegrass) were processed in the same manner to check the accuracy of the isotope measurements. Digested ZnO materials were diluted up to 1:3000. Column yield was checked from the Zn intensities during isotope analysis and was 96.6 ± 18.0 % (mean ± 2SD, n=48) for plant samples and 89.7 ± 5.6 (n=12) for ZnO materials. Three samples with bad yields were rejected and were not included in this study. Zinc fractions were concentration-matched to within 10% and measured for Zn isotopic composition in a High-Resolution Multi-Collector ICP Mass Spectrometer (Neptune, Thermofinnigan). Instrument settings are shown in Supplementary Table 3. The Neptune cup configuration was: L4 (^62^Ni), L2 (^63^Cu), L1 (^64^Zn), Central (^64^Cu), H1 (^66^Zn), H2 (^67^Zn), and H3 (^68^Zn). To correct for mass bias, the sample-standard bracketing procedure as described in^22^ was applied, with AA ETH Zn as bracketing standard and copper (Cu) NIST SRM 976 as the external element. Sample and standard measurements were obtained from 40 cycles of 8-second integrations. Samples were measured 2-3 times on different sessions. Repeatability throughout the three sessions was 0.002%◦ ± 0.035 (2SD; n = 150), calculated from the bracketing standard. Isotope compositions are expressed in the δ notation (Equation 2):

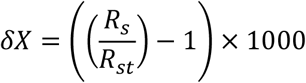

For a given chemical element (X), Rs is the ratio of the heavy isotope to the light isotope of the sample and Rst of the standard. All δ^66^Zn in this study were calculated using the 66 and 64 isotopes and expressed relative to JMC Zn. The δ^66^ZnJMC for AA ETH Zn and ryegrass were 0.27 ± 0.04 (2SD; n = 150), and 0.42 ± 0.06 (2SD; n = 6) respectively, which was in agreement with the literature^23,24^.

### Root anatomy and root cell ultrastructure

Root tips were cut from controls and plants treated with either 1000 mg l^−1^ (Bulk, NP100, NP50) or 10 mg l^−1^ (NW) for 9 weeks. Root tips were rinsed in distilled water and fixed in an ice-cold mixture of 2.5% glutaraldehyde and 2% paraformaldehyde in 0.1M phosphate buffer (pH 7.4), then stored at 4°C. Samples were exposed to vacuum for 1h to remove air bubbles, washed in phosphate buffer, and postfixed and stained with 1% osmium tetroxide and 0.8% potassium ferrocyanide for 1h. Stained samples were washed in distilled water and dehydrated in an acetone series of increasing concentration to achieve 100%. All the fixation steps were carried out at 4°C. Fixed samples were polymerised in epoxy Spurr resin for 48h at 60°C. For light microscopy, 1 μm semi-thin cross-sections were stained with methylene blue and photographed with a light microscope (Olympus 175 CX41, Tokyo, Japan) coupled with a digital camera (Olympus DP70). For transmission electron microscopy (TEM), 70 nm ultrathin sections were cut with a Reichert-Jung Ultracut E ultramicrotome (C. Reichert AG, Vienna, Austria), stained with uranyl acetate and lead citrate, and observed under a Jeol JEM 1010 (Tokyo, Japan) operated at 100 kV. Images were taken with a Gatan Orius camera (Gatan, Pleasanton, USA). For microanalysis, unstained cuts were dried and mounted on titanium grids. Cuts were analysed using a JEOL JEM-2100 LaB6 transmission electron microscope equipped with an Energy Dispersed Analysis of X-ray Spectrometer (EDXS), operating at 200 kV in STEM mode using the dark field detector. The beam size used in this mode was around 15 nm. The spectrometer is an INCA x-sight (Oxford Instruments, Abingdon, UK), with a Si (Li) detector. Micrographs were obtained using a Gatan Orius SC1000 CCD camera with Digital Micrograph Version 1.71.38 software. Map acquisition was accomplished using the INCA Microanalysis Suite version 4.09 software. X-ray maps were obtained by selecting Zn kα1 as the characteristic X-ray peak.

### Evapotranspiration and photosynthetic performance

To calculate the water consumption from each pot (E, in g), the nutrient solution added during each watering was weighed. Chlorophyll content on a leaf area basis was measured with a SPAD-502 portable chlorophyll meter (Minolta, Illinois, USA), as reported elsewhere^25^. Five representative mature leaves of each plant were measured at 2 cm from the base. The leaf gas exchange and chlorophyll fluorescence were determined using a Li-COR 6400 portable photosynthesis system (Li-COR Inc., Lincoln, NE, USA) running OPEN version 4.06. The third fully developed leaf of the healthiest shoot of each plant was measured at approximately 2 cm from the base. Leaves were first dark-adapted for 30 min to measure maximum quantum yield (F_v_/F_m_). The same leaves were then re-acclimated to environmental light until stabilised (up to 45 min) to determine relative quantum yield (F_v_’/F_m_’), quantum yield of photosystem-II photochemistry (ΦPSII)^26^, quantum yield of CO_2_ fixation (ΦCO_2_), the electron transport rate (ETR, μmol m^−2^s^−1^), photochemical (qP) and non-photochemical quenching (qN, NPQ), the light-saturated net CO_2_ assimilation rate (A_s_, μmol CO_2_ m^−2^s^−1^), stomatal conductance to water (g_s_, mol H_2_O m^−2^s^−1^), intercellular CO_2_ concentration (C_i_, μmol CO_2_ mol air^−1^), the transpiration rate (E, mmol H_2_O m^−2^s^−1^), and water vapour pressure deficit of the leaf (VPD, kPa). Measurements were taken under a saturating light (photosynthetic photon flux density of 1200 μmol photons m^−2^ s^−1^), 400 μmol mol^−1^ of CO_2_, and an air temperature of 25.7±0.1°C.

### Carbon and nitrogen isotopic relation in plants

For each plant sample, 0.8-0.9 mg of finely ground dry matter were weighed in tin capsules (Lüdiswiss, Flawil, Switzerland). The total C and N contents were analysed using an Elemental Analyser (EA, Carlo Erba 2100, Milan, Italy), which was interfaced with an Isotope Ratio Mass Spectrometer (IRMS, Thermo-Finnigan Deltaplus Advantage, Bremen, Germany) to analyse the ^13^C/^12^C and ^15^N/^14^N ratios. Results were expressed as δ^13^C and δ^15^N values following Eq. 2, using secondary standards calibrated against Vienna Pee Dee Belemnite calcium carbonate (VPDB) for C, and against N_2_ air for N, respectively. Several certified reference materials were processed in the same manner, at a ratio of one aliquot each per 12 samples. For C, we used IAEA CH7 (measured δ^13^C_VPDB_ −32.1±0.07‰ on n=15, certified −32.2±0.04); IAEA CH6 (measured −10.4±0.08 on n=15, certified 10.5±0.04), and USGS 40 (measured −26.5±0.1 on n=12, certified −26.4±0.04). For N, we analysed IAEA N1 (measured δ^15^N_AIR_ 0.6±2.0‰ on n=14, certified 0.4±0.04), IAEA N2 (measured −20.3±0.8 on n=12, certified 20.3±0.07), IAEA NO_3_ (measured 4.7±0.3 on n=15, certified 4.7±0.2), and USGS 40 (measured −4.5±0.3 on n=11, certified −4.5±0.06).

### Statistical methods

All the statistical analysis was done using R software version 3.4.0 for Windows. Analysis of variance (ANOVA) was performed on each variable based on a two-factor design with interactions. The differences between groups were assessed using paired-t-tests with Bonferroni correction (BF). When data did not meet the assumptions of equal variances or normality, the non-parametric Kruskal-Wallis (KW) ranks test and the Dunn’s test with Benjamini–Hochberg adjustment (Dunn) were used instead. Excel 2016 (Microsoft Office 365 Pro Plus version 1708) and Veusz 1.23.2 were used to create graphs.

## Results

### Characterisation of ZnO sources

Zinc purity was very similar in all four ZnO sources: 94.2% for Bulk, 96.3% for NP100, 90.2% for NP50, and 94.5% for NW (Supplementary Table 4). According to the manufacturer, Zn purity was 99.9%, ~80%, and 91% for Bulk, NP100, and NP50 respectively (no information was given for NW). Besides, 6% Al was reportedly added to NP50 as a dopant but only 2.2% was measured. Traces of other unreported elements were also found: up to 27.2 μg g^−1^ Pb (NP50), 8.5 μg g^−1^ Fe, 4.5 μg g^−1^ Cu, 3.4 μg g^−1^ Ni, 1.2 μg g^−1^ Cd, and 0.7 μg g^−1^ Cr. Average particle length was 215±4 nm (mean±SE, n = 946) for bulk ZnO, with 10% of particles in the nano range. For NP100, the mean particle length was 99±2 nm (n = 956), but 35% of the particles were >100 nm. Similarly, NP50 diameter was 48.7±0.57 nm (n = 467), with 39% of the particles in the 50-100 nm range. Finally, the mean diameter of NW was 59±0.5 nm (n = 223).

### Zinc and Al dissolution and uptake

Zinc concentrations in the growth solution ([Zn]_sol_), roots ([Zn]_root_), and shoots ([Zn]_shoot_) increased with increasing ZnO supply (Fig. 1, Supplementary Table 5), reaching 20x, 231x, and 60x those of the controls at 1000 mg l^−1^, respectively (*P*<0.001). Controls had similar [Zn]_root_ and [Zn]_shoot_ (~30 μg g^−1^). By contrast, plants treated with ZnO accumulated Zn preferentially in the roots (7.6 mg g^−1^ in roots vs. 1.8 mg g^−1^ in shoots at the highest ZnO supply) (Fig. 1B, C). At 100 and 1000 mg l^−1^, NP50 released up to twice as much Zn into solution as bulk and NP100 (*P* =0.019 and 0.011, respectively) (Fig. 1A). [Zn]_shoot_ in plants treated with 1000 mg l^−1^ NP50 reached 3.1 mg g^−1^, twice as much as bulk and NP100 plants (*P* =0.034) (Fig. 1C), and [Zn]_root_ reached 10.4 mg g^−1^ Zn, 60% higher than bulk and 80% higher than NP100 plants (Fig. 1B). The mass balance was calculated to assess the fate of Zn in our system. The proportion of Zn in the solid fraction at the end of the experiment (Zn_solid_) increased with increasing [ZnO] (*P*<0.001) (Fig. 2). At 100 and 1000 mg l^−1^, the majority of Zn (85 - 99 %) remained in the solid fraction. By contrast, the dissolved Zn fraction (Zn_sol_) decreased with increasing ZnO (*P*<0.001). The fraction of Zn stored in roots (Zn_root_) and shoots (Zn_shoot_) gradually increased up to 10 mg l^−1^ and decreased at the two highest concentrations (*P*<0.001). At 100 and 1000 mg l^−1^, Zn_sol_ was the highest in NP50 treatments (9.8 and 2.9%, respectively). The maximum Zn_root_ was 16% and Zn_shoot_ 8%, both at 10 mg l^−1^ NP50.

**Fig. 1.**
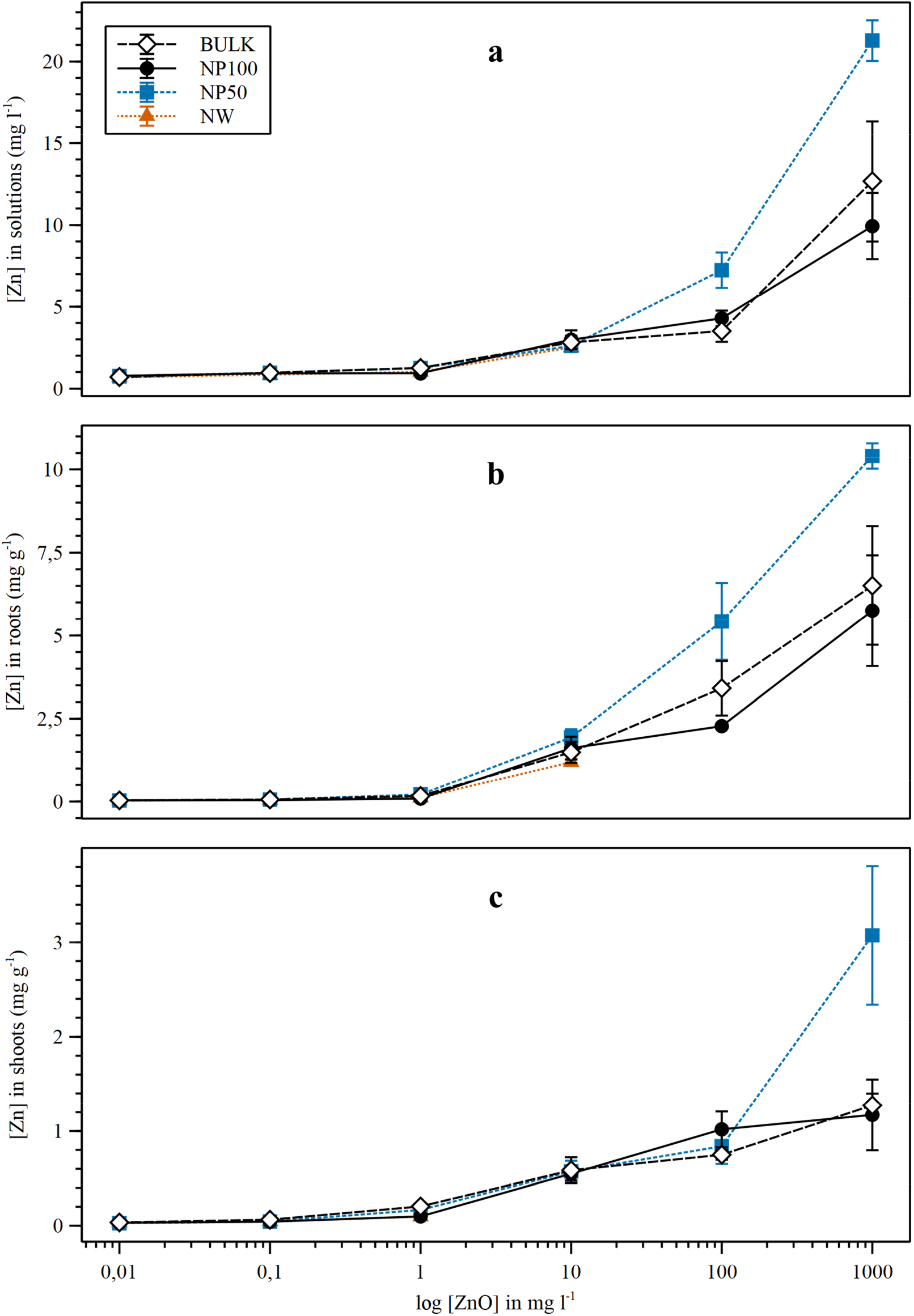
Zinc concentration in nutrient solutions (top), roots (middle), and shoots (bottom). Plants were exposed to four different ZnO sources: micron-size (Bulk), NP < 100 nm (NP100), NP < 50 nm (NP50), and nanowires of 50 nm diameter (NW). Controls (0 mg l^−1^ ZnO) are represented as 0.01 mg l^−1^. Data represent means ±SE, where n = 4. Concentration data are expressed in mg l^−1^ for the growth solutions and in mg g^−1^ for plant samples.

**Fig. 2.**
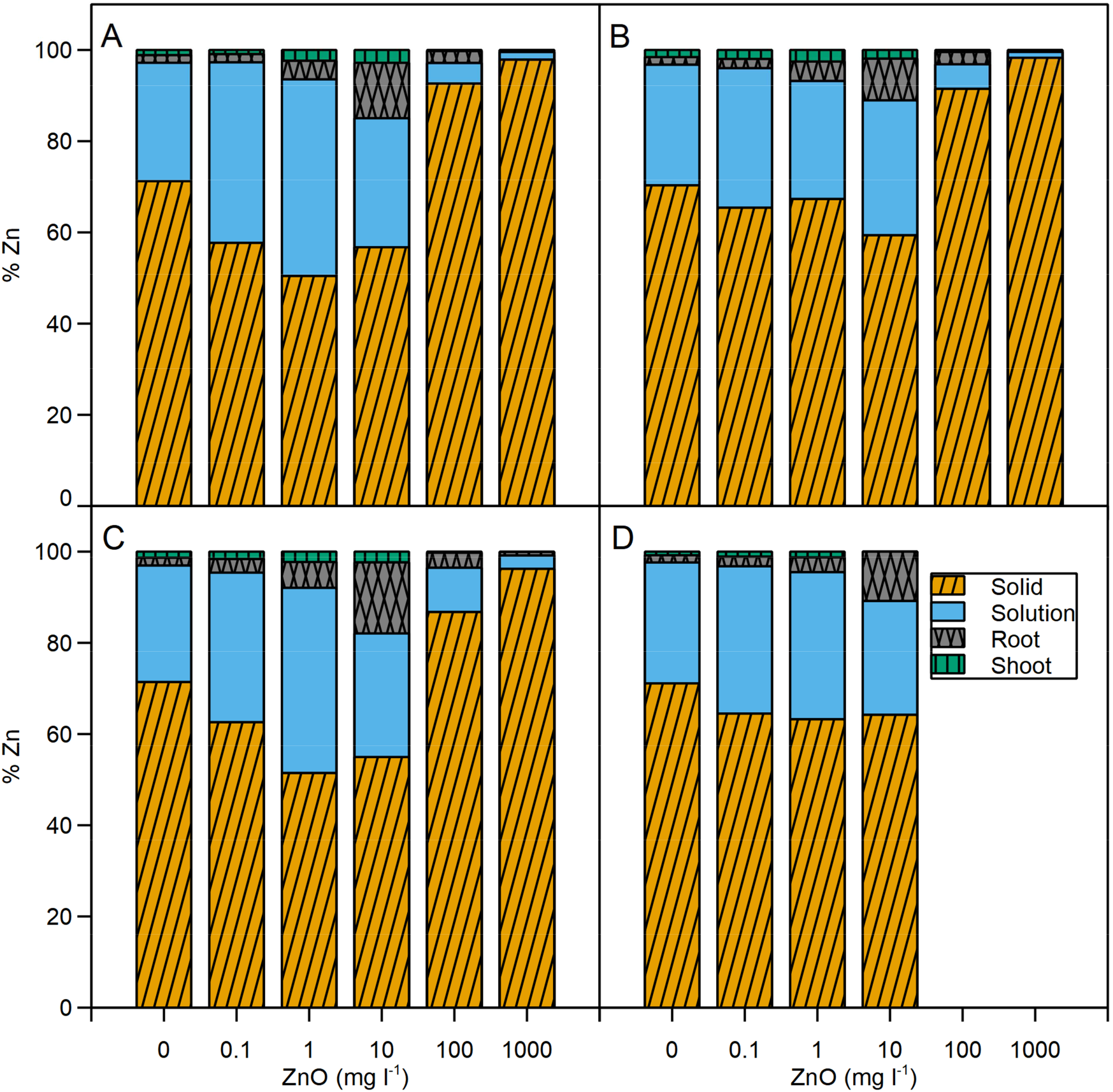
Distribution of Al across the different pools. Plants were treated with four different ZnO sources: micron-size (Bulk), NP < 100 nm (NP100), NP < 50 nm (NP50), and nanowires of 50 nm diameter (NW). Data represent means, where n = 4, expressed as Al % relative to the total Al incorporated into the system from the nutrient solution and ZnO treatments.

To study the uptake of doping elements by plants and the capacity of ENMs to retain metals from solution, Al content and mass balance were investigated. At 1000 mg l^−1^ ZnO (*P*<0.001) (Supplementary Table 5) the [Al]_roots_ was 3.5x higher than the control. The only Al-doped material, NP50, resulted in the highest [Al]_root_ (up to 195 mg g^−1^ at 1000 mg l^−1^, *P*’=0.022). The [Al]shoots increased from the 10 mg l^−1^ ZnO treatments and beyond, with a maximum increase of 30% at 1000 mg l^−1^ ZnO (all *P*<0.001). However, the mass balance calculations indicated that the Al missing from solution mostly ended up in the solid fraction (Alsolid), which increased progressively with [ZnO] (*P*<0.001) (Supplementary Fig. 1). NP50 showed the highest Al_solid_ (*P*<0.001, Supplementary Fig. 1C), starting with 98.9% at 0.1 mg l^−1^. For the rest of the ZnO materials, at 10 mg l^−1^ the Al_solid_ was 42% in Bulk (Supplementary Fig. 1A), 59% in NP100 (Supplementary Fig. 1B), and 78% in NW (Supplementary Fig. 1D). The solution pH progressively increased in pots containing plants, reaching 8.2 by the end of the experiment, while pots without plants had a pH value of 5.5. At week 8, the pH of the 1000 mg l^−1^ solutions was 1 pH unit higher than controls and close to neutrality (*P*<0.001) (Supplementary Table 2). The pH was highest in NP100 treatments and lowest with NW (*P*=0.011). At the end of the experiment, the solution pH was higher for plants under 1000 mg l^−1^ ZnO and controls than for the rest of concentrations, with no difference between sources.

### Zn isotopic fractionation

The δ^66^Zn of all ZnO materials was very similar, on average 0.33 ± 0.04 ‰ (n = 10, Fig. 3, Supplementary Table 6). By contrast, the δ^66^Zn_root_ showed clear differences between treatments (*P*=0.003). Bulk and NP100 plants had the lightest δ^66^Zn_root_ (0.02 and 0.04‰^a^, respectively), followed by NP50 (0.13‰^ab^), and NW and controls (0.35 and 0.38‰^b^, respectively). The δ^66^Zn_shoot_ also differed between ZnO treatments (*P*=0.027, Fig. 3). The shoots of plants treated with bulk ZnO had the lightest δ^66^Zn_shoot_ (−0.61 ‰^a^), while the rest of the ZnO treatments ranged −0.5 to −0.32‰^ab^, and controls had the heaviest δ^66^Zn_shoot_ (0.27‰^b^). The root-to-shoot fractionation (ΔZn_shoot-root_) was very small in controls (−0.08‰), which had no Zn added to the nutrient solution. For the rest of the treatments, ΔZn_shoot-root_ ranged from −0.71 to −0.52‰, with no statistically significant difference among them.

**Fig. 3.**
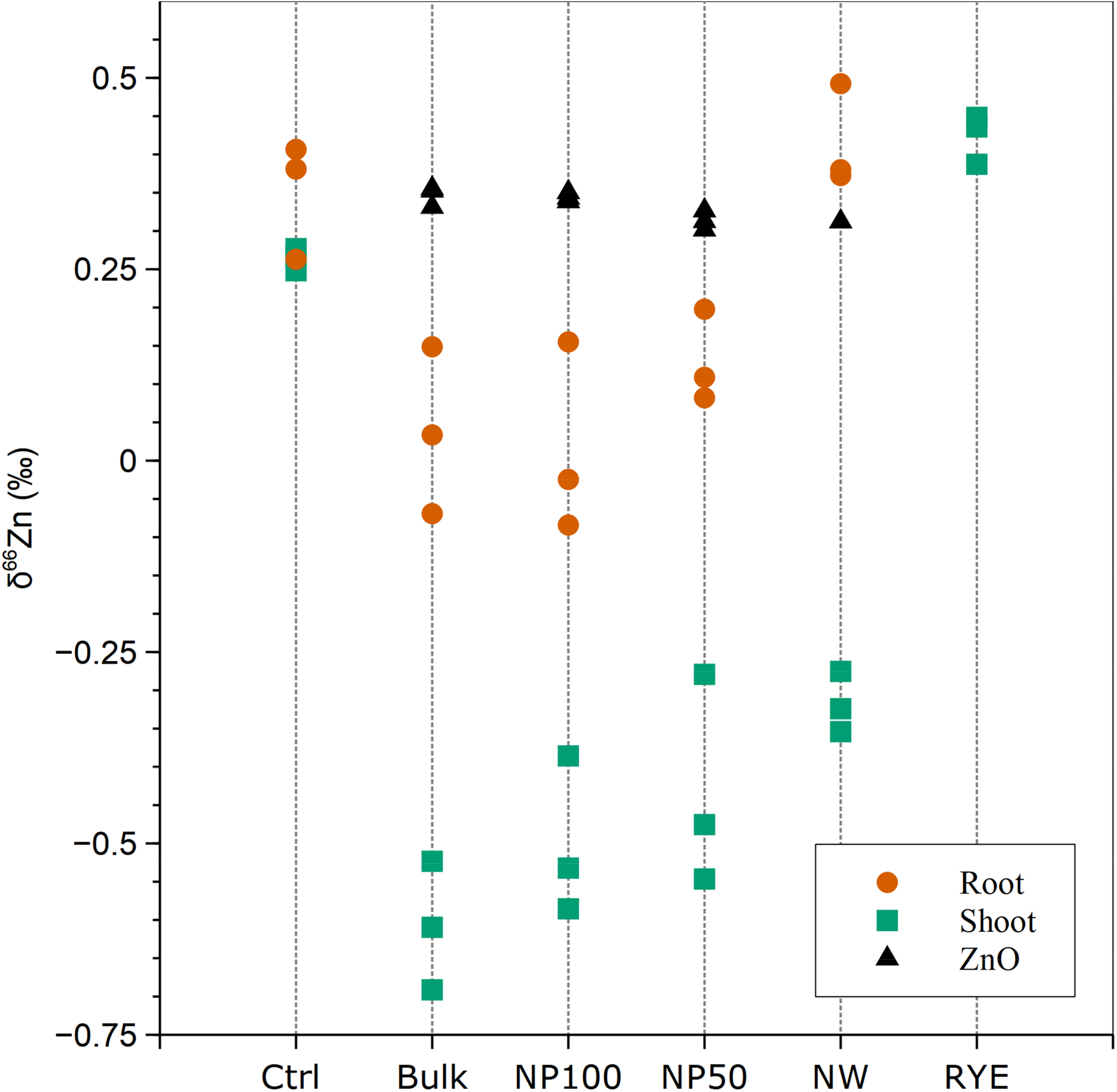
Zinc isotopic composition of ZnO materials, plants, and the BCR-281 (RYE) reference material. Plants were treated with four different ZnO sources at 100 mg l^−1^: micron-size (Bulk), NP < 100 nm (NP100), NP < 50 nm (NP50), and nanowires of 50 nm diameter (NW). Data are expressed relative to JMC Zn.

### Plant growth and evapotranspiration

ZnO caused severe, dose-dependent effects on plant growth and evapotranspiration (ET). Fresh weight (FW) was reduced by up to 74% in whole plants, 66% in roots, and 88% in shoots, whereas FW_root_/ FW_shoot_ was up to 2.6x higher (all *P*<0.001) (Supplementary Table 7). In agreement, dry weight (DW) greatly decreased in response to ZnO, up to 61% in roots and 83% in shoots (both *P*<0.001). The DW_root_/ DW_shoot_ ratio increased up to 2.1x (*P*<0.001). Plant height decreased up to 49% in response to ZnO (*P*<0.001). Root length also decreased with increasing ZnO, with a 61% reduction at 1000 mg l^−1^ (*P*<0.001) (Supplementary Table 7). Mean root length was lowest in NP50 plants (*P* =0.005). The rest of the growth parameters measured did not show any significant differences between sources. The ET decreased up to 59% with ZnO (*P*<0.001). ET from plants at 1000 mg l^−1^ was 1078±41 g of solution (n = 14), which was very similar to pots without plants (992±24, n = 5), evidencing a strong inhibition of plant transpiration. Significant effects on growth and ET started from 1 mg l^−1^ (DW_shoot_, DW_root_/ DW_shoot_, ET), 10 mg l^−1^ (all FW, DW_root_, height), or 100 mg l^−1^ (root length), but the trend was often present at 0.1 mg l^−1^. The δ^13^C increased in shoots at [ZnO] ≥100 mg l^−1^ (*P*<0.001) (Supplementary Table 8). A similar trend was observed in roots but it did not attain significance.

### Photosynthetic performance

The chlorophyll content of mature leaves decreased in response to all ZnO materials and was 25% lower than controls at 1000 mg l^−1^ (*P*<0.001, Supplementary Table 9). Accordingly, photosynthetic efficiency was severely affected. A_s_ and ΦCO_2_ decreased from 10 mg l^−1^ and dropped by 79% and 75% respectively at 1000 mg l^−1^ (*P*<0.001) (Fig. 4, Supplementary Table 9). The ΦPSII, qP, and ETR also decreased dramatically, up to 80% at 1000 mg l^−1^ (all *P*<0.001). By contrast, F_v_/F_m_ had only a minor reduction (6%) (*P*=0.002), whereas NPQ increased by 4.4x at 1000 mg l^−1^ (*P*=0.005). The ΦCO_2_, F_v_/F_m_, ΦPSII, qP, and ETR were lower in plants under the Bulk treatment, followed by the NP50 treatment (*P*=0.046, 0.038, 0.002, 0.014, and 0.002, respectively, Fig. 4, Supplementary Table 9). Gas exchange was also affected by ZnO: g_s_ and E decreased up to 78% and 72%, respectively (both *P*<0.001), while VPD was 20% higher than controls at 1000 mg l^−1^ (*P*=0.002). Finally, F_v_’/F_m_’ was not affected by ZnO.

**Fig. 4.**
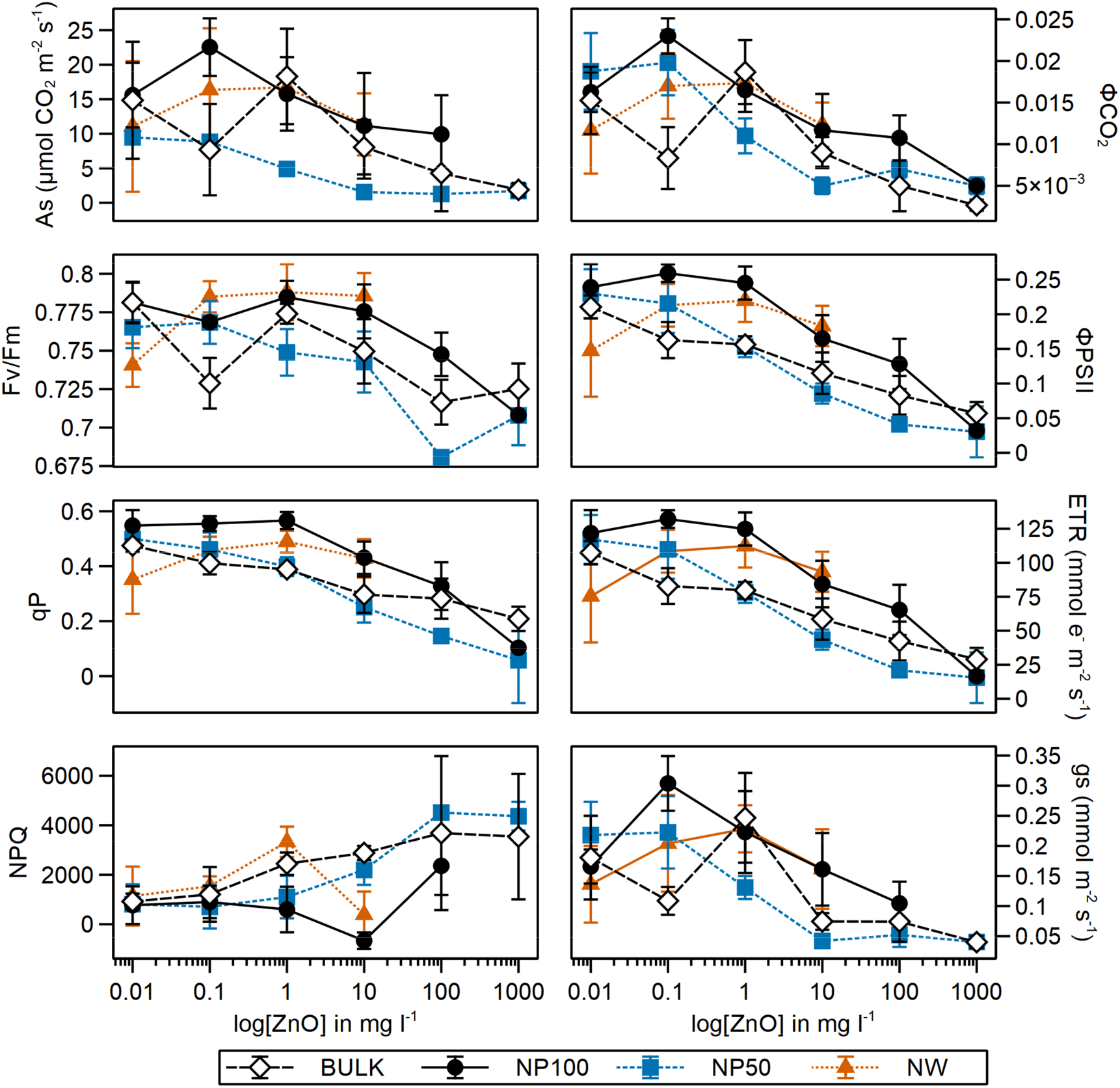
Photosynthetic performance under ZnO stress. Plants were treated with four different ZnO sources: micron-size (Bulk), NP < 100 nm (NP100), NP < 50 nm (NP50), and nanowires of 50 nm diameter (NW). Data represent means ±SE, where n = 4. The variables F_v_/F_m_ (maximum quantum yield of PSII photochemistry), ΦPSII (quantum yield of PSII electron transport, qP (photochemical quenching), and NPQ (non-photochemical quenching) are dimensionless.

### Root anatomy and ultrastructure

Initial light microscope exploration revealed extensive damage caused by ZnO to the roots: detachment of the epidermis, disorganisation and thickening of cell walls, and vacuolisation and death of root cells, which is indicative of increased aerenchyma formation (Supplementary Fig. 2). Plants treated with NW showed the most severe effects, with generalised apoptosis of the cortex cells that made further characterisation difficult. On the 100Kev TEM, the root epidermis showed significant loss of cell wall material in the Bulk, NP100, and NP50 treatments (Supplementary Fig. 3). The epidermal cells in the Bulk and NP50 treatments exhibited loss of turgor, protoplasm shrinkage, and accumulation of electron-dense granules in the vacuoles. In the cortex, cell-wall thickening, protoplasm shrinkage, and a high number of starch granules or amyloplasts were observed in all three treatments (Supplementary Fig. 4). Additionally, electron-dense precipitates accumulated in the intercellular spaces (Supplementary Fig. 5).

Microanalysis confirmed the presence of Zn in various locations: i) a few granules on the root surface in NP50 and NP100 that are morphologically consistent with NPs (Fig.5); ii) large amorphous precipitates between cell walls and plasma membrane in the cortex of NP50, NP100, and Bulk plants (in order of abundance, Fig.6–7), where most of Zn accumulated; and iii) small vacuoles containing Zn, P, and sometimes Ca in the rhizodermis and cortex of NP50 plants (Fig.6).

**Fig. 5.**
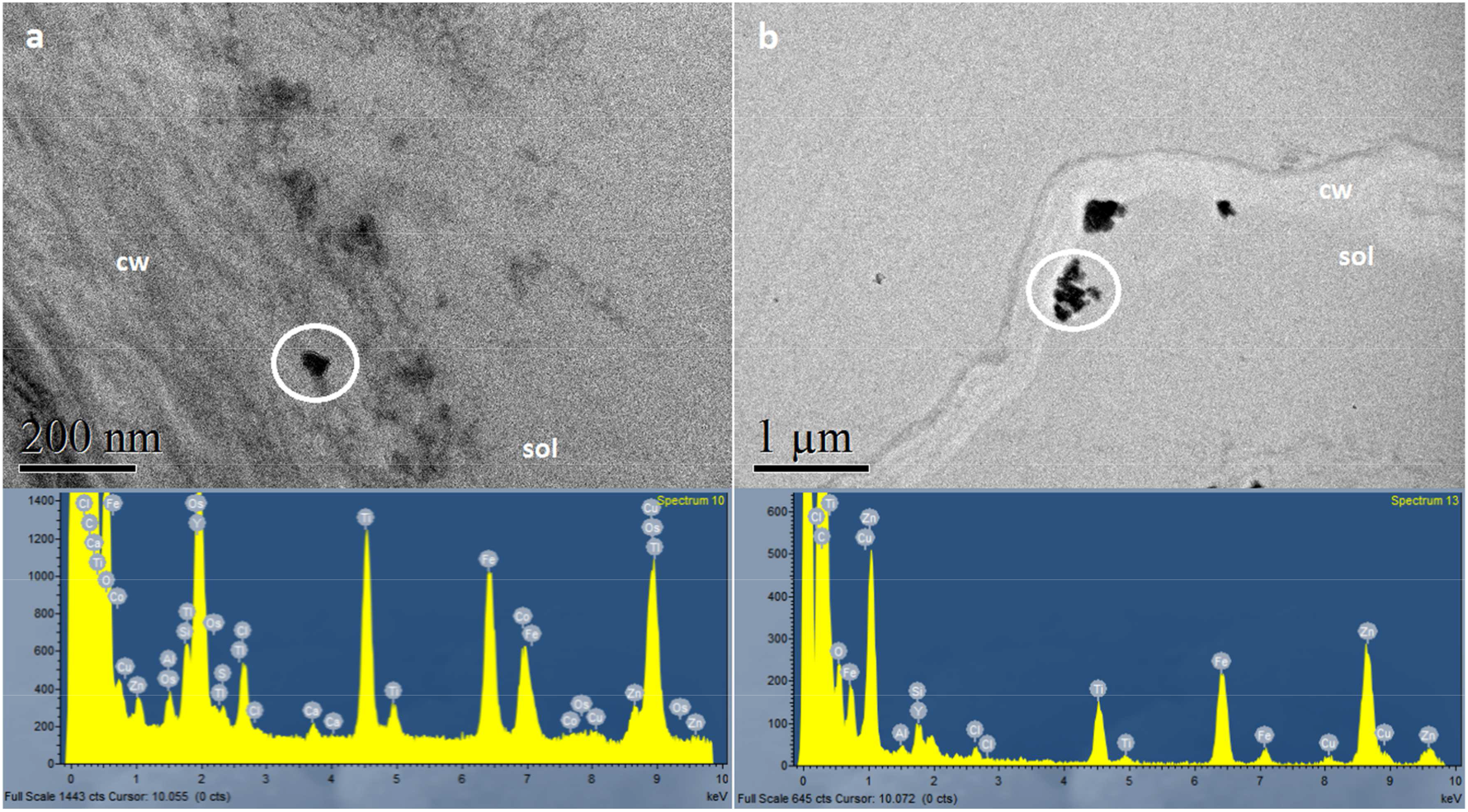
ZnO-NPs on the root surface. Roots in (a) were treated with NP < 50 nm (NP50), while roots in (b) were treated with NP < 100 nm (NP100). Spectra represent X-Ray spectrometry analysis of the circled areas. Cell wall = cw; exterior solution = sol.

**Fig. 6.**
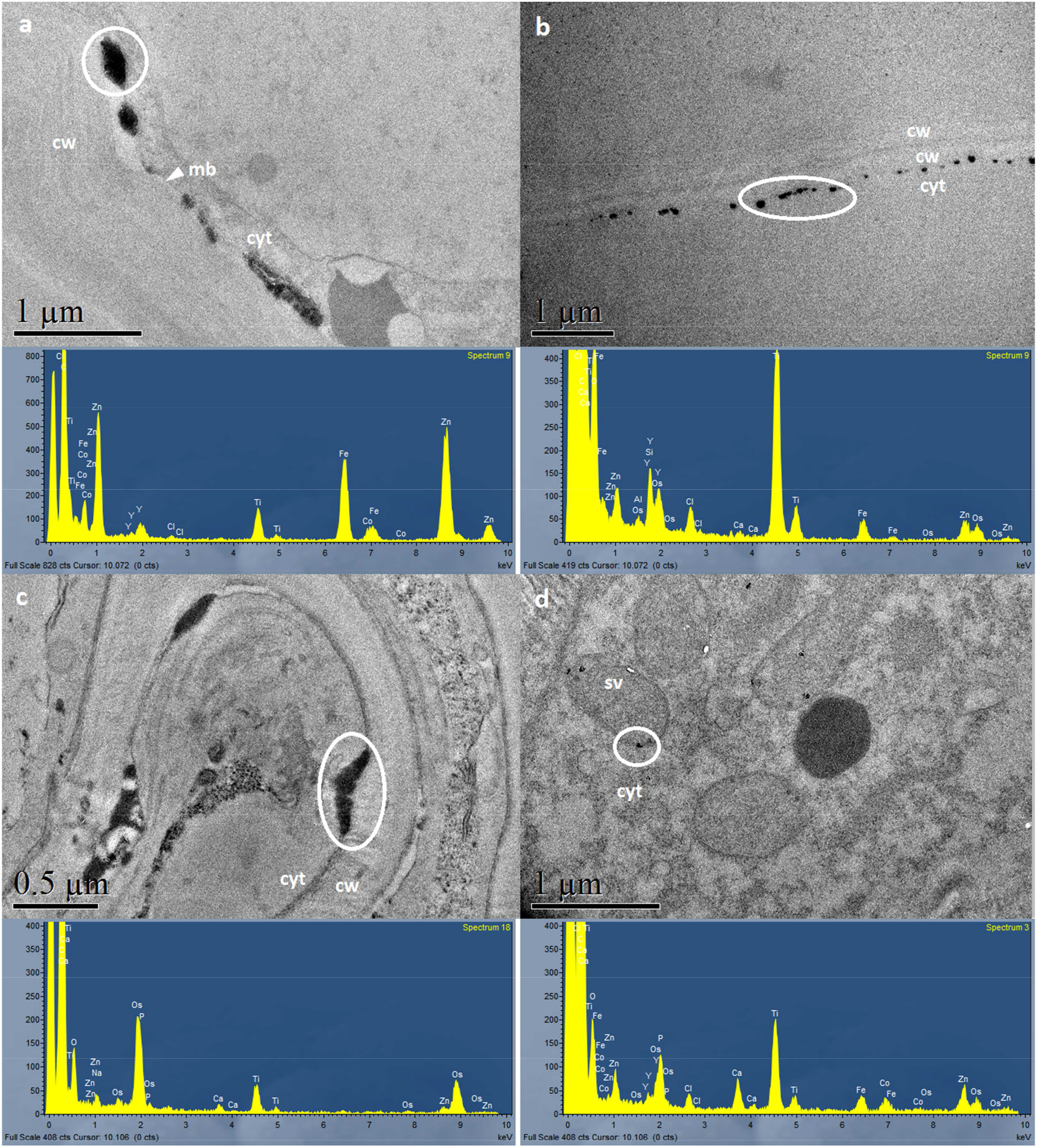
Zinc deposits in the root cortex. (a), (b), and (c) show Zn precipitates between the cell walls and plasma membranes in NP50, NP100, and Bulk plants, respectively. (d) shows Zn sequestration in small vacuoles in NP50 plants. Spectra represent X-Ray spectrometry analysis of the circled areas. Cell wall = cw; cytoplasm = cyt; plasma membrane = mb; small vacuole = sv.

**Fig. 7.**
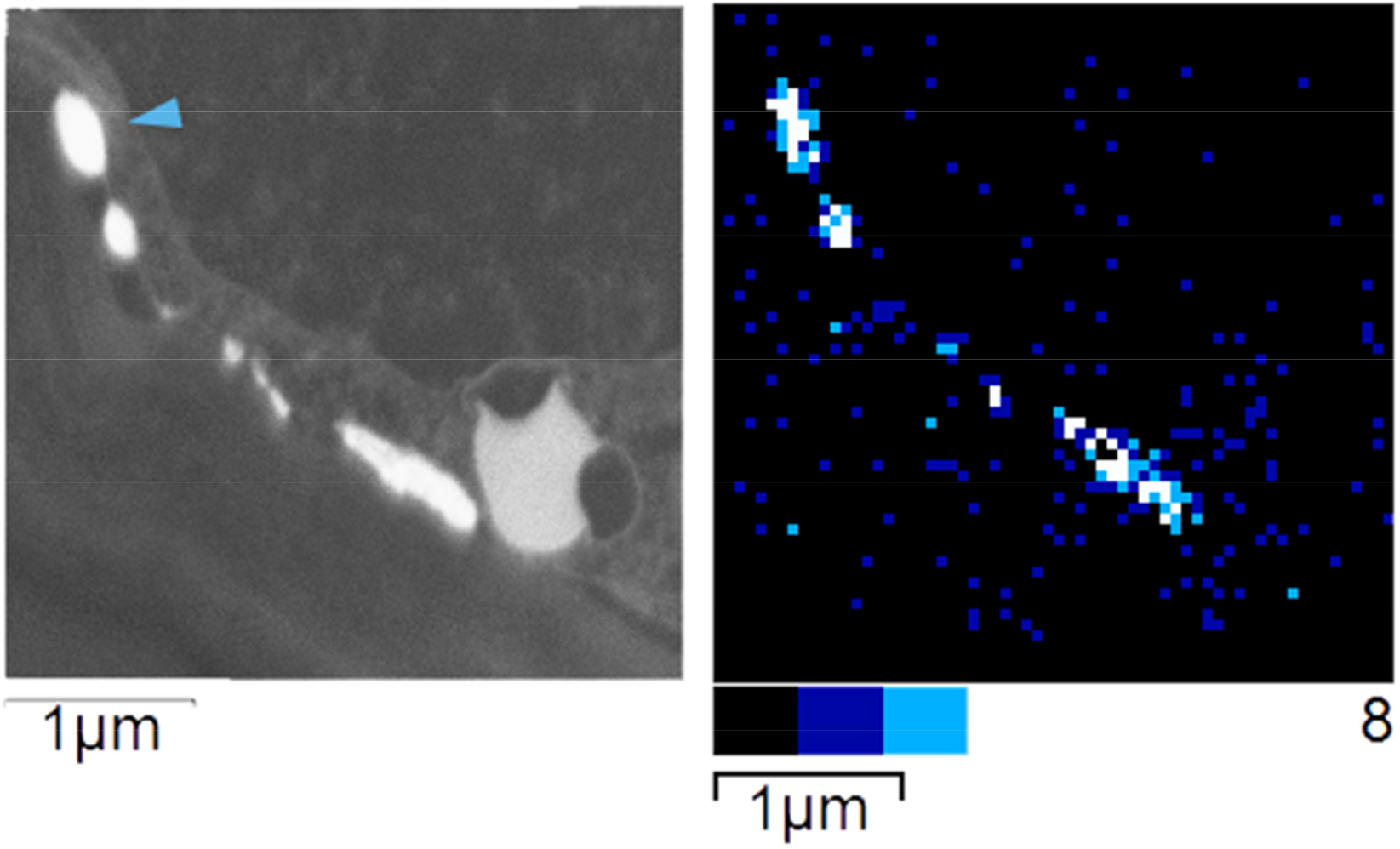
Dark-field micrograph and elemental map of the same region as Fig.6a. The Zn precipitates were located between the cell walls and plasma membranes in the NP50 root cortex. The colour scale indicates the intensity of the Zn signal in counts, from 0 (black) to 8 (white).

### Nutrient content and distribution

Nitrogen content changed in response to ZnO, with a N_shoot_ increase of up to 24% (*P*<0.001, Supplementary Table 8). The Cshoot/Nshoot ratio decreased progressively in response to increasing ZnO, reaching a 17% reduction at 1000 mg l^−1^ (*P*<0.001). ZnO slightly increased C_root_ (~5% at 1000 mg l^−1^, *P* =0.023) and decreased C_shoot_ (~3% at 100 mg l^−1^, *P* =0.013). The C_root_/C_shoot_ ratio increased progressively with increasing [ZnO], up to 7% (*P* =0.023). In the roots, [Mn] and [Fe] greatly increased in response to ZnO, up to 79% and 99%, respectively, at 1000 mg l^−1^ (*P*=0.012 and 0.004, Supplementary Table 9). Conversely, [K] showed a progressive decrease that reached 41% at 1000 mg l^−1^ ZnO (*P*<0.001). Phosphorus in the roots increased in response to low ZnO, then decreased up to 25% at 1000 mg l^−1^ (*P*<0.001). In the shoots, [P] was higher than controls at 10 mg l^−1^ ZnO and above, with a peak 34% increase at 100 mg l^−1^ (*P*<0.001). Besides, both [S] and [K] increased in response to 10-100 mg l^−1^ ZnO, attaining levels 32% and 14% higher, respectively, than controls at 100 mg l^−1^ (*P*<0.001 and *P*=0.020). Copper in shoots increased at 0.1 and 1 mg l^−1^ ZnO, then gradually decreased across treatments, reaching a 30% reduction at 1000 mg l^−1^ (*P*<0.001). Manganese in shoots progressively decreased in response to ZnO and was 66% lower than controls at the highest ZnO level (*P*<0.001). However, [Mn]_shoot_ was lower in the NP50 and NP100 treatments compared to the other ZnO sources (*P*<0.001).

## Discussion

### Route of plant exposure to ZnO-ENMs

Our results show that ZnO-ENMs dissolve slowly in the chosen experimental conditions and that the majority was still solid at the end of the experiment (Fig. 2). Dissolved Zn reached only 4.3-7.2 mg l^−1^ in 100 mg l^−1^ ZnO-ENM treatments (Fig.1). This range closely agrees with dissolution experiments by Reed and co-workers, where 100 mg l^−1^ ZnO-NP suspended in water for 60 days released 2.2-7.4 mg l^−1^ into solution^27^. Smaller NPs are known to dissolve more easily due to their larger surface area^28^, which explains why [Zn]_sol_ was higher in the NP50 treatment than in the NP100 and bulk treatments (Fig.1). Solution pH was ~6 initially but gradually increased up to 7.5-8.2 in the presence of plants. This pH range is typical of wetland waters^29^. In our study, the proportion of undissolved Zn and Al increased with increasing ZnO concentrations and solution pH. Accordingly, removal of Zn, Cd, Cu, Hg, Ni, and Pb from solution by ZnO-ENMs has been reported to increase with pH and increasing sorbent mass^30,31^. The effect of pH on metal removal is explained by the increased deprotonation of the surface and attraction between negative hydroxyl sites and cations, while the effect of the sorbent mass is due to the increased number of binding sites. Aluminium is frequently added as a dopant to ZnO-NPs to enhance their electrical and optical properties^32^, so the Al mass balance in our system can serve as a proxy for the fate of doping elements. In our study, most of the Al ended up in the solid fraction, which increased with increasing ZnO, while the dissolved Al fraction decreased in the presence of ZnO (Supplementary Fig.1). However, [Al] was higher in plants grown at high [ZnO], especially for Al-doped NP50. This is consistent with the “Trojan horse effect” hypothesis, which states that ENMs might bind to other chemicals in solution and enhance their uptake by plants, leading to increased phytotoxicity. Aluminium is toxic to plants, with a reduction in root elongation as the main symptom^33^. The increased uptake of Zn and Al could explain the stronger reduction in root length in NP50 plants as compared with other treatments (Supplementary Table 6). Additionally, the capacity of ZnO-ENMs for metal removal from solution might have affected the bioavailability of essential nutrients, as seen from the changes in nutrient content.

In all ZnO treatments, roots showed intense vesicularisation at the plasma membrane, but the vesicles did not contain either ZnO-ENMs or Zn. Instead, Zn was found mostly in amorphous precipitates outside the plasma membrane, and in NP50 plants alone, in small vacuoles containing Zn, Ca, and P (Fig.6). Very few ZnO-ENMs were found attached to the root surface and none were detected in the cytoplasm (Fig.5). These observations indicate that ZnO-ENMs dissolve and the roots take up Zn^2+^, which they mostly immobilise as Zn precipitates in the apoplast and Zn phosphates in small vacuoles. There is abundant evidence in the literature that plant roots sequester excess Zn as precipitates in the intercellular spaces or store Zn in vacuoles bound to phosphates, organic acids or phytochelatins^34–39^. Plants exposed to ZnO-ENMs also accumulate Zn in roots in the form of amorphous phosphates or phytates^12,14,40^. In our study, Zn and P only co-occurred in the vacuoles of NP50 plants (Fig.6). The higher [Zn] achieved in the roots of NP50 plants might have activated vacuolar sequestration. Alternatively, NPs of this size might have been captured by endocytosis and then dissolved inside the vacuole and sequestered as phosphates. The endocytosis of entire 50 nm ZnO-NPs has been reported in hydroponically grown rice^10^. However, our results do not support this explanation because no NPs were detected inside the cells in our study.

The Zn isotope data support the predominant uptake of Zn^2+^, followed by sequestration of excess Zn as precipitates and complexes in root cells. The main causes of Zn isotopic fractionation in plants are Zn speciation, compartmentalisation, and the activity of membrane transporters^20^. Divalent Zn uptake by low-affinity transporters at the plasma membrane favours the light isotopes, which diffuse faster across membranes thanks to their smaller size^20,41^. Hence, the relative accumulation of light Zn isotopes in the Bulk, NP100, and NP50 roots compared to ZnO (Fig. 3) can be explained by ZnO dissolution followed by Zn^2+^ uptake mediated by membrane transporters. However, NP50 roots were to some extent enriched in heavy isotopes compared to NP100 and bulk roots, although this tendency was not significant. This can be attributed to a greater number and size of Zn precipitates in the intercellular spaces of NP50 roots, which also contained more Zn relative to other treatments, and to the presence of Zn-rich vacuoles in the cortex (Fig.6–7). The formation of Zn precipitates and complexes in roots during the plant response to excess Zn^2+^ left an isotopically lighter pool of Zn^2+^ for transportation to the shoot and favoured the accumulation of heavy isotopes in the root^42–44^. Nevertheless, the δ^66^Zn of NP50 roots remained lighter than the starting ZnO because this isotopic composition was still dominated by the initial transfer through the membrane, which favours light isotopes. NW roots were very damaged and had lost their structural coherence, which explains why these roots had an isotopic composition that equalled the ZnO of the NW. Further, the shoots became equally depleted in heavy isotopes in all treatments relative to the roots: ΔZn_shoot-root_ was −0.64±0.22, - 0.52±0.25, −0.57±0.12, and −0.71±0.01 for Bulk, NP100, NP50, and NW50, respectively. The magnitude and direction of ΔZn_shoot_-root are in the same range as previously observed in *P. australis* in response to ZnCl2 toxicity^44^. Depletion of heavy Zn isotopes during Zn transport to the shoot is attributed to preferential sequestration of heavy isotopes in the root, the activity the membrane transporters during Zn loading into the xylem, and bulk flow^44–46^. Conversely, there is no isotopic fractionation when Zn is transported in complexes up the shoot due to the bigger mass of the complexes^47^. The same likely applies to ZnO-ENMs. Hence, our isotope data indicate that Zn was predominantly transported up the shoot as Zn^2+^, regardless of the ZnO source, and that plants roots are an effective barrier against ZnO-ENM transport up the shoot, which is in agreement with our TEM observations. If ZnO-ENM had been transported through the plant without dissolution, then no isotopic fractionation would have been detected during Zn uptake and transport in the plants. ZnO-NPs in the apoplast reportedly can reach the symplast by endocytosis^11^ or through the discontinuity of Casparian bands^6,14^. However, ZnO-NPs in our study were found on the surface of the root but not in the apoplast, indicating that the rhizodermis was an effective barrier. Similarly, recent research shows that ZnO-NPs are unlikely to reach the shoots unless roots are damaged^13^. Finally, the fractionation of Zn isotopes in control plants is consistent with the plants’ response to Zn-deficiency, which results in Zn uptake and transport to the shoots in the form of Zn complexes, with root exudates enriched in heavy isotopes^48^. The δ^66^Zn of all the ZnO materials was remarkably similar (0.33‰) and the value was consistent with reported ranges for both ZnO-ENMs (−0.31 to 0.28‰)^49^ and natural Zn minerals like hydrozincite (0-0.30‰)^50^.

### Phytotoxicity

Chronic exposure to high levels of ZnO-ENMs decreased growth, chlorophyll content, and photosynthetic assimilation in *P. australis* (Fig.4, Supplementary Tables 6-7). Biomass allocation, nutrient uptake and distribution, and fractionation of C isotopes were also altered by ZnO-ENMs (Supplementary Tables 8-9). Similar effects have been previously described in aquatic macrophytes treated with toxic levels of ZnO-ENMs^4–6,8,10,51,52^, Zn^2+^ (reviewed by ^20^), and other metals ^53–56^. In our previous research on *P. australis*, exposure to 2 mM ZnCl_2_ (131 mg l^−1^ Zn in solution) for 40 days caused a height reduction of 25%^44^. Song and coworkers^4^ reported a ~50% reduction in height after exposure to 100 mg l^−1^ ZnO NP50 (only 4.5 mg l^−1^ Zn in solution) for 35 days^4^. Even though [Zn^2+^] was lower in the latter study, the effects were more severe and attributed to direct root contact with ZnO-NPs. Accordingly, we report a height decrease of 43% after three months at 100 mg l^−1^ ZnO (3.5-7.2 mg l^−1^ in solution), irrespective of particle size. The magnitude of the effect paralleled the Song *et al*. study despite the longer exposure time and larger particle sizes of the Bulk, NP100, and NW treatments. To sum up, ZnO is more toxic to *P. australis* than ZnCl2. The total ZnO added to the system is the most important factor to explain the impact of ZnO-ENMs on plant growth, rather than [Zn^2+^] in solution and particle size. The toxicity cannot be attributed to the osmotic effect of the increased [Zn^2+^]. The conductance (EC) did not increase with increasing ZnO, which remained mostly undissolved. This clearly demonstrates that undissolved ZnO plays a key role in ENM phytotoxicity. To correctly assess the environmental risks of ZnO-ENMs, field studies should include the solid ZnO fraction in sediments, rather than just focus on ZnO-ENM levels in the water. However, particle size was relevant for ZnO-ENM impacts on root growth and [Mn]_shoot_ due to the greater dissolution of smaller NPs.

The lower photosynthetic quantum yield ofates can be explained by stomatal closure and root malfunction leading to water deficit, higher δ^13^C, lower chlorophyll content, and nutritional imbalance. Of particular interest were the changes in Mn content, which greatly decreased in shoots in a dose-dependent response to [ZnO] and particle size (Supplementary Table 9). It is unlikely that this decrease in [Mn]_shoot_ is caused by Mn removal from solution by ENMs. The removal efficiency is low for Mn^57^ and the [Mn]_root_ was high, proving that Mn uptake was not impaired. We propose that excess Zn^2+^ might inhibit or compete for Mn^2+^ transporters involved in Mn root-to-shoot transport, like AtZIP1 and ATZIP2. These carriers transport Zn and Mn, are expressed in the root vasculature, and mediate Mn radial movement towards the xylem and xylem loading^58^. Manganese has a key role as a catalyst of the oxygen-evolving complex of PSII and its deficiency greatly reduces photosynthesis^59^. Gas exchange and chlorophyll fluorescence data taken at the same time confirm lower electron transport efficiency. E and gs decreased, indicating that the stomatal opening was strongly inhibited. The reduced gas exchange could lower A_s_ by restricting CO_2_ availability. The parallel increase in δ^13^C of shoot and root dry matter also supports this conclusion. However, Ci was not affected in our study. Helophytes take up CO_2_ dissolved in water by the roots, and it diffuses towards the photosynthetic tissues via the aerenchyma^60^. This alternative source of CO_2_ can contribute significantly to Ci^61,62^. The small 5% reduction in F_v_/F_m_ in dark-adapted leaves and the unaltered F_v_’/F_m_’ showed that PSII was mostly functional. By contrast, the ΦPSII, qP, ΦCO_2_, and ETR were severely reduced in light-adapted leaves, while qN and NPQ increased. Hence, we attribute the reduced ΦCO_2_ and As to decreased stomatal conductance and lower efficiency of electron transport downstream due to an Mn deficiency that caused PSII to become easily saturated by light. Reduced chlorophyll content, As, gs, E, Ci, F_v_/F_m_, qP, and ETR coupled with increased NPQ have been reported in terrestrial plants exposed to toxic levels of ZnO-ENMs^63,64^. In agreement, the aquatic plants *Azolla filiculoides* and *Lemna minor* have shown severely reduced growth, chlorophyll content and F_v_/F_m_ after exposure to ZnO-ENMs at pH 4.5-5.5^5,10,65^. Remarkably, growth and F_v_/F_m_ in *L. minor* were not equally reduced at pH 8, due to the lower NP dissolution^65^. In our study, *P. australis* progressively basified the growth media, which reached pH~8 at the end of the experiment. This high final pH could explain the limited effect of ZnO on F_v_/F_m_. The δ^13^C_shoot_ was within −26.7 to −30.7 ‰, which is in agreement with values previously reported for C3 helophytes^53,66^. In C3 plants, diffusion of atmospheric CO_2_ through the stomatal pore and C fixation by RuBisCO favour ^12^C^67^. During stomatal closure, RuBisCO continues consuming intercellular CO_2_ and the δ C_shoot_ increases as ^12^C is depleted^68^. In our study, δ C_shoot_ and δ^13^C_root_ increased (~1‰) in response to high ZnO. This, in the context of reduced transpiration, is generally caused by Ci exhaustion^67^. However, our gas exchange data do not indicate Ci depletion. While *P. australis* mainly assimilates atmospheric C, it can also take up CO_2_ dissolved in water by the roots^60^. Plants that assimilate more CO_2_ by this route show higher δ^13^C, as seen when comparing submerged aquatic plants relative to helophytes^66^. The assimilation of a larger proportion of CO_2_ from the roots during stomatal closure can explain a higher δ^13^C while maintaining Ci levels. In agreement, we observed increased root aerenchyma development in response to excess Zn. We found no previous record of δ^13^C in plants exposed to toxic levels of ZnO-ENMs, although *Iris pseudacorus* grown in 200 mg l^−1^ ZnCl_2_ showed a similar increase of δ^13^C_shoot_^53^.

Our experiment shows that ZnO-ENMs are toxic from 1 mg l^−1^. Effects at this concentration include reduced growth, allocation of new biomass to the roots and rhizomes instead of the photosynthetic tissues, and restriction of transpiration. These levels are high compared to the few existing records of ZnO-ENMs in surface waters: there are up to 1.84 μg l^−1^ ZnO NMs in the surface waters of Singapore^69^ and just 5 ng l^−1^ in Canada^70^. However, recent modelling studies estimate ZnO-ENM concentrations of 10^−6^ to 1 μg l^−1^ in surface waters and 11-32 mg kg^− 1^ in sediments^71–73^. Current levels predicted for the sediments would cause substantial toxic effects on *P. australis*. Besides, the said models have important limitations, and the real concentrations are likely to be locally higher and to increase in coming years^74^.

## Conclusion

Emerging nano-pollutants like ZnO-ENM are increasingly discharged into surface waters and they accumulate in sediments. This is a cause of major concern because of our limited understanding of the mechanisms of ZnO-ENM uptake and toxicity in wetland plants, which play key ecological roles in aquatic ecosystems. Besides, there is an ongoing controversy about the capacity of ENMs to enter plant roots and be transported to the shoots. Our research concludes that ZnO-ENM dissolve before uptake by plant roots and that they are not transported to the shoots. Nanoparticles smaller than 50 nm dissolve faster than the larger particles and are more toxic to plants. Exposure to ZnO-ENM takes place mostly through the uptake of dissolved Zn^2+^, but also through direct contact with the root surface. The mechanisms of action of ZnO-ENM adsorbed onto the root surface require further investigation. In *Phragmites australis*, ZnO-ENM reduce shoot and root growth, transpiration, and photosynthetic rate, while inducing changes in nutrient uptake and distribution. This could be detrimental to the environmental health of aquatic ecosystems and their capacity to sequester carbon. Wetlands are one of the major carbon sinks globally. Pollutants that compromise the biomass production of wetland plants can increase global carbon emissions, of vital importance in the current context of climate change. In conclusion, current and future ZnO-ENM levels in sediments could pose a significant risk for aquatic plants and ecosystems. Our research also evidences the importance of considering the undissolved fraction, exposure time, and NP size to correctly evaluate the environmental risk of nanomaterials.

## Acknowledgements

This project has received funding from the European Union’s Horizon 2020 research and innovation programme under the Marie Skłodowska-Curie grant agreement NANOREM No 704957. Franck Poitrasson is funded by the French Centre National de la Recherche Scientifique. We are grateful for the technical assistance of Joan Martorell and Luís López with TEM microanalysis; Manuel Henry and Jérôme Chmeleff with Zn isotope analysis; and Josep Mata, Marta Pintó, and Xavier García with greenhouse tasks and photosynthesis measurements. We thank Sheela Paramjothy for her help with maintaining the experiment.

