## Supplemental Info for "Stable Zn isotopes reveal the uptake and toxicity of zinc oxide engineered nanomaterials in *Phragmites australis*"

Supplementary Table 1. Composition of the nutrient solution.

| | Stock solution | | Half-strength | |
|---|---|---|---|---|
| $KNO_3$ | 101.1 | $g\ l^{-1}$ | 303.0 | $mg\ l^{-1}$ |
| $Ca(NO_3)_2 \cdot 4H_2O$ | 236.2 | $g\ l^{-1}$ | 472.0 | $mg\ l^{-1}$ |
| $NH_4H_2PO_4$ | 115.1 | $g\ l^{-1}$ | 115.0 | $mg\ l^{-1}$ |
| $MgSO_4 \cdot 7H_2O$ | 246.5 | $g\ l^{-1}$ | 123.0 | $mg\ l^{-1}$ |
| NaFeEDTA (13.7-18.7% Fe) | 30.00 | $g\ l^{-1}$ | 15.00 | $mg\ l^{-1}$ |
| KCl | 1.864 | $g\ l^{-1}$ | 1.864 | $mg\ l^{-1}$ |
| $H_3BO_3$ | 0.773 | $g\ l^{-1}$ | 773.0 | $\mu g\ l^{-1}$ |
| $MnSO_4 \cdot H_2O$ | 0.169 | $g\ l^{-1}$ | 169.0 | $\mu g\ l^{-1}$ |
| $H_2SO_4$ (98%) | 54.00 | $\mu l\ l^{-1}$ | 54.00 | $\mu l\ ml^{-1}$ |
| $CuSO_4 \cdot 5H_2O$ | 62.00 | $mg\ l^{-1}$ | 62.00 | $\mu g\ l^{-1}$ |
| $H_2MoO_4$(85% $MoO_3$) | 40.00 | $mg\ l^{-1}$ | 40.00 | $\mu g\ l^{-1}$ |
| pH | | | 5.86 | |
| EC (µS) | | | 1243 | |

Modified from[72].

Supplementary Table 2. Conductivity and pH of nutrient solutions.

| | | [ZnO] (mg $l^{-1}$) | | | | | | | Source | | | | |
|---|---|---|---|---|---|---|---|---|---|---|---|---|---|
| | No plant | 0 | 0.1 | 1 | 10 | 100 | 1000 | $\chi^2$ | Bulk | NP100 | NP50 | NW | $\chi^2$ |
| pH | | | | | | | | | | | | | |
| 8 weeks | 4.2±0.1 | 5.8±0.2$^{ab}$ | 5.6±0.1$^{a}$ | 5.6±0.2$^{ab}$ | 5.8±0.1$^{ab}$ | 6.2±0.1$^{b}$ | 6.8±0.1$^{c}$ | 38.9*** | 5.9±0.1$^{ab}$ | 6.3±0.1$^{b}$ | 5.9±0.2$^{ab}$ | 5.6±0.2$^{a}$ | 9.5* |
| 12 weeks | 5.5±0.4 | 8.1±0.1$^{c}$ | 7.6±0.1$^{ab}$ | 7.6±0.1$^{ab}$ | 7.5±0.05$^{a}$ | 7.9±0.1$^{bc}$ | 8.2±0.1$^{c}$ | 28.2*** | 7.9±0.1 | 7.9±0.1 | 7.8±0.1 | 7.7±0.1 | 3.1 |
| EC (µS) | | 43.2±0.2$^{ab}$ | 44.0±0.1$^{b}$ | 42.9±0.2$^{a}$ | 43.4±0.8$^{ab}$ | 41.8±1$^{a}$ | 42.5±0.6$^{a}$ | 14.5* | 5.9±0.2$^{a}$ | 6.3±0.8$^{ab}$ | 5.9±1$^{a}$ | 5.6±0.6$^{a}$ | 3.1 |

Plants were grown in four different ZnO sources: micron-size (Bulk), nanoparticles < 100 nm (NP100), nanoparticles < 50 nm (NP50), and nanowires of 50 nm diameter (NW). Each source was provided at six concentrations (0, 0.1, 1, 10, 100, and 1000 mg $l^{-1}$) except NW (only up to 10 mg $l^{-1}$). Data represent means ±SE, where n = 4 and df= 5 for the ZnO concentration factor and df = 3 for the source factor. Concentration data are expressed in mg $l^{-1}$ for the growth solutions and in µg $g^{-1}$ for plant samples. Different letters indicate statistically significant groups according to Dunn's test with Benjamini-Hochberg correction. The Chi-square value ($\chi^2$) corresponds to the Kruskal-Wallis rank-sum test and is indicated as significant at $P < 0.05$ (*), $P < 0.01$ (**), or $P < 0.001$(***).

Supplementary Table 3. Neptune multicollector typical operational settings.

| | |
|---|---|
| Extraction [V]: | -2000 |
| Focus [V]: | -666.3 |
| SourceQuad1 [V]: | 248.7 |
| Rot-Quad1 [V]: | 0 |
| Foc-Quad1 [V]: | -17.6 |
| Rot-Quad2 [V]: | 29.5 |
| Source Offset [V]: | -6 |
| Matsuda Plate [V]: | 17.5 |
| Cool Gas [l/min]: | 15.1 |
| Aux Gas [l/min]: | 0.9 |
| Sample Gas [l/min]: | 1 |
| Add Gas [l/min]: | 0 |
| Org Gas [l/min]: | 0 |
| Operation Power [W]: | 1308 |
| X-Pos [mm]: | 2.5 |
| Y-Pos [mm]: | 1.5 |
| Z-Pos [mm]: | 2.9 |
| Ampl.-Temp [°C]: | 47.23 |
| Fore Vacuum [mbar]: | 4.17E-04 |
| High Vacuum [mbar]: | 3.91E-07 |
| Ion Getter-Press [mbar]: | 2.48E-08 |

Supplementary Table 4. Elemental composition of ZnO materials.

| | Source | | | |
|---|---|---|---|---|
| | Bulk | NP100 | NP50 | NW |
| [Al] µg $g^{-1}$ | 0.6±0.3 | 0.9±0.3 | 22400±2500 | 6.2±1.9 |
| [Cd] µg $g^{-1}$ | 1.2±0.5 | 0.9±0.4 | 1.1±0.1 | 0.0±0.0 |
| [Cr] µg $g^{-1}$ | 0.1±0.0 | 0.1±0.0 | 0.3±0.0 | 0.7±0.1 |
| [Cu] µg $g^{-1}$ | 4.5±1.7 | 3.1±1.4 | 3.2±0.5 | 0.4±0.0 |
| [Fe] µg $g^{-1}$ | 1.3±0.3 | 0.9±0.4 | 8.4±1.0 | 8.5±1.1 |
| [Ni] µg $g^{-1}$ | 0.1±0.0 | 0.1±0.0 | 0.5±0.1 | 3.4±0.3 |
| [Pb] µg $g^{-1}$ | 3.7±1.4 | 2.7±1.2 | 27.2±3.9 | 0.2±0.0 |
| [Zn] mg $g^{-1}$ | 756.8±7.2 | 773.9±1.6 | 724.9±15.2 | 759.6±15.1 |

ZnO materials were: micron-size (Bulk), nanoparticles < 100 nm (NP100), nanoparticles < 50 nm (NP50), and nanowires of 50 nm diameter (NW). Data represent means ±SE, n =3. Data are expressed in mg $g^{-1}$ for Zn and in µg $g^{-1}$ for the rest of the elements.

Supplementary Table 5. Zinc and aluminium concentrations in solutions and plants.

| | [ZnO] (mg $l^{-1}$) | | | | | | | Source | | | | |
|---|---|---|---|---|---|---|---|---|---|---|---|---|
| | 0 | 0.1 | 1 | 10 | 100 | 1000 | $\chi^2$ | Bulk | NP100 | NP50 | NW | $\chi^2$ |
| [Zn] | | | | | | | | | | | | |
| solution | 0.73±0.02$^{a}$ | 0.92±0.03$^{ab}$ | 1.12±0.05$^{b}$ | 2.71±0.17$^{c}$ | 5.01±0.6$^{cd}$ | 14.76±1.8$^{d}$ | 84.6*** | 3.53±1.0 | 3.57±0.8 | 5.92±1.5 | 1.31±0.2 | 6.6 |
| root | 33.1±3.0$^{a}$ | 50.7±3.4$^{ab}$ | 150±14.0$^{b}$ | 1537±133$^{bc}$ | 3705±581$^{cd}$ | 7630±932$^{d}$ | 86.3*** | 1869±555 | 1799±537 | 3073±791 | 376±125 | 6.7 |
| shoot | 31.5±3.6$^{a}$ | 48.5±4.4$^{a}$ | 143±15$^{b}$ | 601±46$^{c}$ | 868±89$^{c}$ | 1880±370$^{c}$ | 82.6*** | 467±94 | 512±123 | 821±252 | 211±69 | 4.1 |
| [Al] | | | | | | | | | | | | |
| solution | 0.14±0.01$^{bc}$ | 0.17±0.01$^{c}$ | 0.15±0.01$^{c}$ | 0.11±0.02$^{ab}$ | 0.07±0.01$^{a}$ | 0.10±0.01$^{a}$ | 39.0*** | 0.13±0.01 | 0.13±0.01 | 0.12±0.01 | 0.14±0.01 | 3.0 |
| root | 31.4±3.5$^{a}$ | 23.8±3.1$^{a}$ | 39.4±4.7$^{ab}$ | 32.9±4.9$^{a}$ | 32.0±5.3$^{a}$ | 110.4±28.6$^{b}$ | 19.2** | 35.1±4.2$^{ab}$ | 36.2±8.9$^{a}$ | 67.4±15$^{b}$ | 28.6±3.1$^{a}$ | 9.6* |
| shoot | 19.7±3.4$^{ab}$ | 14.7±1.6$^{a}$ | 15.4±1.6$^{a}$ | 25.4±2.1$^{c}$ | 21.9±2.7$^{abc}$ | 26.1±3.5$^{bc}$ | 25.1*** | 22.2±2.5 | 16.3±1.4 | 21.6±2.4 | 20.8±2.3 | 5.0 |
| [Al]/[Zn] | | | | | | | | | | | | |
| solution | 0.19±0.01$^{d}$ | 0.19±0.01$^{cd}$ | 0.14±0.01$^{c}$ | 0.044±0.006$^{b}$ | 0.017±0.003$^{ab}$ | 0.009±0.002$^{a}$ | 80.6*** | 0.1±0.01 | 0.1±0.02 | 0.09±0.02 | 0.14±0.01 | 4.4 |
| root | 0.66±0.06$^{e}$ | 0.32±0.04$^{de}$ | 0.14±0.03$^{cd}$ | 0.046±0.005$^{bc}$ | 0.026±0.003$^{ab}$ | 0.016±0.002$^{a}$ | 81.6*** | 0.23±0.06 | 0.19±0.05 | 0.17±0.04 | 0.33±0.06 | 6.9 |
| shoot | 1.0±0.1$^{c}$ | 0.46±0.04$^{bc}$ | 0.31±0.05$^{b}$ | 0.022±0.003$^{a}$ | 0.0090±0.0008$^{a}$ | 0.015±0.004$^{a}$ | 81.6*** | 0.32±0.08 | 0.26±0.06 | 0.32±0.1 | 0.44±0.09 | 4.6 |

Plants were grown in four different ZnO sources: micron-size (Bulk), nanoparticles < 100 nm (NP100), nanoparticles < 50 nm (NP50), and nanowires of 50 nm diameter (NW). Each source was provided at six concentrations (0, 0.1, 1, 10, 100, and 1000 mg $l^{-1}$) except NW (only up to 10 mg $l^{-1}$). Data represent means ±SE, where n = 4 and df= 5 for the ZnO concentration factor and df = 3 for the source factor. Concentration data are expressed in mg $l^{-1}$ for the growth solutions and in µg $g^{-1}$ for plant samples. Different letters indicate statistically significant groups according to Dunn's test with Benjamini-Hochberg correction. The Chi-square value ($\chi^2$) corresponds to the Kruskal-Wallis rank-sum test and is indicated as significant at $P < 0.05$ (*), $P < 0.01$ (**), or $P < 0.001$(***).

Supplementary Table 6. $\delta^{66}Zn_{JMC}$ of plant tissues and ZnO sources.

| | Control | Bulk | NP100 | NP50 | NW | test | value |
|---|---|---|---|---|---|---|---|
| $\delta^{66}Zn_{shoot}$ (‰) | $0.27\pm0.01^{b}$ | $-0.61\pm0.05^{a}$ | $-0.50\pm0.06^{ab}$ | $-0.44\pm0.08^{ab}$ | $-0.32\pm0.02^{ab}$ | KW | 11.0* |
| $\delta^{66}Zn_{root}$ (‰) | $0.35\pm0.05^{b}$ | $0.04\pm0.06^{a}$ | $0.02\pm0.07^{a}$ | $0.13\pm0.04^{ab}$ | $0.38\pm0.01^{b}$ | AOV | 9.1** |
| $\delta^{66}Zn_{ZnO}$ (‰) | | $0.35\pm0.01^{a}$ | $0.34\pm0.00^{a}$ | $0.31\pm0.01^{a}$ | $0.31^{a}$ | KW | 6.7 |

Plants were grown in four different ZnO sources: micron-size (Bulk), nanoparticles < 100 nm (NP100), nanoparticles < 50 nm (NP50), and nanowires of 50 nm diameter (NW). Each source was provided at 100 mg $l^{-1}$ except NW (10 mg $l^{-1}$). Data represent means ±SE, where n =3 and df= 4. Different letters indicate statistically significant groups according to either paired-t-tests with Bonferroni correction (ANOVA, AOV) or Dunn's test with Benjamini-Hochberg correction (Kruskal-Wallis, KW). The F-value (AOV) or Chi-square value (KW) is indicated as significant at P < 0.05 (*), P < 0.01 (**), or P < 0.001(***).

Supplementary Table 7. Plant growth and evapotranspiration (ET) in response to ZnO.

| | [ZnO] (mg $l^{-1}$) | | | | | | | | Source | | | | | |
|---|---|---|---|---|---|---|---|---|---|---|---|---|---|---|
| | 0 | 0.1 | 1 | 10 | 100 | 1000 | test | value | Bulk | NP100 | NP50 | NW | test | value |
| $Fw_{plant}$ | 56.8±8.2$^{d}$ | 53.4±9.7$^{d}$ | 34.4±5.0$^{cd}$ | 26.9±3.1$^{bc}$ | 18.7±2.4$^{ab}$ | 15.0±0.9$^{a}$ | KW | 37.3*** | 27.1±3.8 | 41.8±6.5 | 38.2±7.5 | 36.0±4.0 | KW | 5.3 |
| $FW_{root}$ | 36.4±4.5$^{c}$ | 34.9±5.6$^{bc}$ | 25.5±3.6$^{bc}$ | 20.4±2.1$^{ab}$ | 14.7±1.8$^{a}$ | 12.5±0.7$^{a}$ | KW | 32.6*** | 19.9±2.3 | 28.4±3.5 | 25.6±4.6 | 26.8±2.6 | KW | 6.3 |
| $FW_{shoot}$ | 15.5±3.6$^{c}$ | 12.2±3.5$^{bc}$ | 6.4±1.2$^{abc}$ | 4.9±0.9$^{ab}$ | 2.9±0.6$^{ab}$ | 1.8±0.3$^{a}$ | AOV | 35.6*** | 5.0±1.2 | 10.7±2.9 | 8.4±2.2 | 6.4±1.1 | AOV | 4.3 |
| $FW_{root}/FW_{shoot}$ | 3.4±0.4$^{a}$ | 4.6±0.6$^{ab}$ | 5.2±0.7$^{abc}$ | 6.1±1.1$^{bc}$ | 6.4±1.0$^{bc}$ | 8.8±1.0$^{c}$ | KW | 23.0*** | 5.6±0.5 | 4.7±0.6 | 6.3±1.0 | 5.9±0.8 | KW | 3.4 |
| $DW_{root}$ | 5.3±0.6$^{d}$ | 4.7±0.6$^{cd}$ | 3.8±0.5$^{bcd}$ | 2.9±0.4$^{abc}$ | 2.5±0.4$^{ab}$ | 2.0±0.2$^{a}$ | KW | 26.5*** | 3.0±0.4 | 4.1±0.4 | 3.6±0.6 | 4.0±0.4 | KW | 6.1 |
| $DW_{shoot}$ | 4.2±0.8$^{d}$ | 3.2±0.8$^{cd}$ | 1.9±0.4$^{bc}$ | 1.6±0.3$^{bc}$ | 1.1±0.1$^{ab}$ | 0.7±0.1$^{a}$ | KW | 36.4*** | 1.6±0.3 | 2.8±0.7 | 2.4±0.5 | 2.0±0.3 | KW | 3.4 |
| $Dw_{root}/DW_{shoot}$ | 1.6±0.2$^{a}$ | 2.2±0.3$^{ab}$ | 2.3±0.2$^{b}$ | 2.0±0.1$^{ab}$ | 2.4±0.2$^{bc}$ | 3.4±0.3$^{c}$ | KW | 23.2*** | 2.3±0.2 | 2.3±0.2 | 2.2±0.2 | 2.4±0.2 | KW | 1.1 |
| Height | 56.2±3.4$^{d}$ | 50.2±4.4$^{cd}$ | 46.3±2.9$^{cd}$ | 42.7±2.9$^{bc}$ | 32.3±1.7$^{ab}$ | 28.5±1.9$^{a}$ | KW | 37.0*** | 40.6±2.9 | 45.7±3.0 | 45.3±3.7 | 43.1±2.7 | KW | 1.6 |
| Root length | 38.6±2.5$^{c}$ | 38.0±2.2$^{c}$ | 33.6±2.7$^{bc}$ | 30.2±2.8$^{bc}$ | 23.9±2.7$^{ab}$ | 14.8±1.3$^{a}$ | AOV | 12.0*** | 28.5±2.3$^{ab}$ | 31.4±2.2$^{ab}$ | 27.6±2.7$^{a}$ | 37.7±2.8$^{b}$ | AOV | 4.6** |
| ET | 2656±266$^{d}$ | 2336±282$^{cd}$ | 1769±192$^{bc}$ | 1683±128$^{bc}$ | 1373±82$^{ab}$ | 1078±41$^{a}$ | KW | 40.4*** | 1594±139 | 2093±220 | 1851±223 | 1962±152 | KW | 5.9 |

FW = fresh weigth; DW = dry weight; and ET = Evapotranspiration, all in g. Height and root length in cm. Plants were grown in four different ZnO sources: micron-size (Bulk), nanoparticles < 100 nm (NP100), nanoparticles < 50 nm (NP50), and nanowires of 50 nm diameter (NW). Each source was provided at six concentrations (0, 0.1, 1, 10, 100, and 1000 mg $l^{-1}$) except NW (only up to 10 mg $l^{-1}$). Data represent means ±SE, where n = 4 and df= 5 for the ZnO concentration factor and df = 3 for the source factor. Different letters indicate statistically significant groups according to either paired-t-tests with Bonferroni correction (ANOVA, AOV) or Dunn's test with Benjamini-Hochberg correction (Kruskal-Wallis, KW). The F-value (AOV) or Chi-square value (KW) is indicated as significant at $P < 0.05$ (*), $P < 0.01$ (**), or $P < 0.001$(***).

Supplementary Table 8. Carbon and Nitrogen content and isotopic composition of plant samples.

| | [ZnO] (mg l$^{-1}$) | | | | | | | | Source | | | | | |
|---|---|---|---|---|---|---|---|---|---|---|---|---|---|---|
| | 0 | 0.1 | 1 | 10 | 100 | 1000 | test | value | Bulk | NP100 | NP50 | NW | test | value |
| C | | | | | | | | | | | | | | |
| root | 42.1±0.2$^{a}$ | 43.4±0.2$^{b}$ | 44.1±0.9$^{ab}$ | 44.4±0.7$^{b}$ | 43.3±0.9$^{ab}$ | 44.3±0.7$^{b}$ | K | 13.1* | 44.2±0.6 | 44.3±0.6 | 42.8±0.4 | 42.8±0.3 | K | 4.9 |
| shoot | 43.2±0.2$^{ab}$ | 44.0±0.1$^{b}$ | 42.9±0.2$^{a}$ | 43.4±0.8$^{ab}$ | 41.8±1$^{a}$ | 42.5±0.6$^{a}$ | K | 14.5* | 42.8±0.4 | 43.6±0.4 | 42.4±0.4 | 43.5±0.4 | K | 3.1 |
| root/shoot | 0.98±0.01$^{a}$ | 0.99±0.005$^{ab}$ | 1.03±0.02$^{bc}$ | 1.03±0.02$^{bc}$ | 1.04±0.03$^{bc}$ | 1.05±0.03$^{c}$ | K | 13.0* | 1.03±0.02 | 1.02±0.02 | 1.01±0.01 | 0.99±0.01 | K | 1.8 |
| N | | | | | | | | | | | | | | |
| root | 2.8±0.1$^{a}$ | 3.1±0.1$^{b}$ | 3.0±0.1$^{a}$ | 3.1±0.2$^{ab}$ | 2.7±0.1$^{a}$ | 2.8±0.2$^{a}$ | K | 1.5** | 2.9±0.1 | 2.9±0.1 | 2.9±0.1 | 3.0±0.1 | K | 0.4 |
| shoot | 3.3±0.1$^{a}$ | 3.7±0.1$^{b}$ | 3.6±0.1$^{b}$ | 3.7±0.2$^{b}$ | 3.8±0.2$^{b}$ | 4.1±0.2$^{b}$ | K | 23.5*** | 3.7±0.1 | 3.7±0.2 | 3.7±0.1 | 3.6±0.1 | K | 0.4 |
| root/shoot | 0.85±0.02 | 0.83±0.02 | 0.83±0.03 | 0.85±0.06 | 0.74±0.06 | 0.71±0.07 | K | 7.4 | 0.81±0.03 | 0.82±0.05 | 0.78±0.03 | 0.83±0.03 | K | 2.1 |
| C/N | | | | | | | | | | | | | | |
| root | 15.3±0.3$^{ab}$ | 14.2±0.3$^{a}$ | 14.9±0.5$^{ab}$ | 15.2±0.9$^{ab}$ | 16.1±0.6$^{ab}$ | 16.8±1.0$^{b}$ | K | 13.2* | 15.5±0.5 | 15.7±0.7 | 15.2±0.4 | 14.7±0.5 | K | 2.1 |
| shoot | 13.2±0.3$^{c}$ | 11.9±0.2$^{b}$ | 11.8±0.2$^{b}$ | 12.0±0.5$^{b}$ | 11.2±0.7$^{ab}$ | 11.0±0.9$^{a}$ | K | 27.1*** | 11.9±0.4 | 12.1±0.5 | 11.6±0.4 | 12.2±0.3 | K | 1.3 |
| $\delta^{13}C$ | | | | | | | | | | | | | | |
| root | -28.0±0.1 | -27.9±0.2 | -27.7±0.1 | -27.8±0.2 | -27.6±0.2 | -27.3±0.2 | K | 6.9 | -27.9±0.1 | -27.7±0.2 | -27.7±0.1 | -27.6±0.1 | K | 3.0 |
| shoot | -28.9±0.2$^{a}$ | -29.3±0.2$^{a}$ | -28.9±0.1$^{a}$ | -29.0±0.2$^{a}$ | -28.6±0.3$^{ab}$ | -28.0±0.2$^{b}$ | A | 6.4*** | -28.8±0.2 | -28.8±0.2 | -28.9±0.1 | -28.8±0.2 | A | 0.2 |
| root/shoot | 0.97±0.003 | 0.95±0.004 | 0.96±0.01 | 0.96±0.01 | 0.97±0.01 | 0.98±0.01 | K | 5.2 | 0.97±0.01 | 0.96±0.01 | 0.96±0.004 | 0.96±0.01 | K | 1.3 |
| $\delta^{15}N$ | | | | | | | | | | | | | | |
| root | -0.87±0.33 | -0.66±0.29 | -0.21±0.37 | -0.36±0.31 | -0.48±0.40 | 0.13±0.18 | A | 6.5 | -0.24±0.27 | -0.56±0.26 | -0.53±0.26 | -0.34±0.28 | A | 1.5 |
| shoot | 0.35±0.20 | 0.46±0.21 | 0.34±0.26 | 0.39±0.19 | 0.62±0.39 | 1.06±0.2 | A | 2.2 | 0.61±0.18 | 0.12±0.24 | 0.68±0.17 | 0.68±0.14 | A | 2.5 |
| root/shoot | 3.41±1.9 | 0.56±2.4 | -3.74±3.0 | 0.38±1.5 | -0.21±0.4 | 0.39±0.3 | K | 3.8 | -0.7±2.03 | 0.92±1.33 | 0.75±1.11 | -0.7±2.59 | K | 2.4 |

Plants were grown in four different ZnO sources: non-nano (Bulk), nanoparticles < 100 nm (NP100), nanoparticles < 50 nm (NP50), and nanowires of 50 nm diameter (NW). Each source was provided at six concentrations (0, 0.1, 1, 10, 100, and 1000 mg $l^{-1}$) except NW (only up to 10 mg $l^{-1}$). Data represent means ±SE, where n = 4 and df= 5 for the ZnO concentration factor and df = 3 for the source factor. Concentration data are expressed in mg $l^{-1}$ for the growth solutions, in % for the C and N content of plant samples, and in ‰ for $\delta^{13}C$ and $\delta^{15}N$ of plant samples. Different letters indicate statistically significant groups according to either paired-t-tests with Bonferroni correction (ANOVA, A) or Dunn's test with Benjamini-Hochberg correction (Kruskal-Wallis, K). The F-value (ANOVA) or Chi-square value (Kruskal-Wallis) is indicated as significant at $P < 0.05$ (*), $P < 0.01$ (**), or $P < 0.001$(***).

Supplementary Table 9. Photosynthetic performance

| | [ZnO] (mg $l^{-1}$) | | | | | | | |
|---|---|---|---|---|---|---|---|---|
| | 0 | 0.1 | 1 | 10 | 100 | 1000 | value | test |
| SPAD | 40.3±1.6$^{bc}$ | 43.8±1.3$^{c}$ | 42.2±1.6$^{c}$ | 38.4±1.4$^{bc}$ | 34.0±2.3$^{ab}$ | 30.8±1.1$^{a}$ | 35.9*** | KW |
| Fv/Fm | 0.77±0.007$^{b}$ | 0.77±0.007$^{b}$ | 0.77±0.008$^{b}$ | 0.76±0.01$^{b}$ | 0.73±0.012$^{a}$ | 0.72±0.01$^{a}$ | 19.1** | KW |
| ΦPSII | 0.21±0.019$^{d}$ | 0.22±0.017$^{d}$ | 0.19±0.014$^{cd}$ | 0.14±0.017$^{bc}$ | 0.10±0.022$^{ab}$ | 0.04±0.016$^{a}$ | 11.7*** | ANOVA |
| qP | 0.48±0.033$^{c}$ | 0.47±0.026$^{c}$ | 0.46±0.025$^{bc}$ | 0.36±0.037$^{abc}$ | 0.29±0.052$^{ab}$ | 0.13±0.068$^{a}$ | 27.1*** | KW |
| ETR | 107404±9611$^{cd}$ | 109922±8578$^{d}$ | 97758±7282$^{cd}$ | 71174±8749$^{bc}$ | 51170±11389$^{ab}$ | 21403±8172$^{a}$ | 11.7*** | ANOVA |
| qN | 361±194$^{a}$ | 430±195$^{a}$ | 469±208$^{ab}$ | 404±211$^{ab}$ | 401±263$^{b}$ | 205±204$^{b}$ | 13.8* | KW |
| NPQ | 894±343$^{a}$ | 1083±347$^{a}$ | 1681±462$^{ab}$ | 1257±502$^{ab}$ | 3130±1196$^{b}$ | 3890±997$^{b}$ | 16.6** | KW |
| Fv'/Fm' | 2.1±0.06 | 2.1±0.07 | 2.3±0.15 | 2.5±0.23 | 2.3±0.09 | 2.5±0.21 | 9.7 | KW |
| $\Phi CO_2$ | 0.016±0.002$^{bc}$ | 0.018±0.002$^{c}$ | 0.015±0.001$^{bc}$ | 0.01±0.001$^{ab}$ | 0.008±0.002$^{a}$ | 0.004±0.001$^{a}$ | 25.8*** | KW |
| A | 15.1±2.0b$^{c}$ | 17.1±2.1$^{c}$ | 14.7±1.4$^{bc}$ | 9.0±1.5$^{ab}$ | 7.3±1.7$^{a}$ | 3.1±0.6$^{a}$ | 26.1*** | KW |
| $g_s$ | 0.18±0.026$^{bc}$ | 0.22±0.033$^{c}$ | 0.2±0.026$^{bc}$ | 0.11±0.025$^{abc}$ | 0.08±0.019$^{ab}$ | 0.04±0.006$^{a}$ | 27.8*** | KW |
| E | 4.3±0.58$^{bc}$ | 4.8±0.61$^{c}$ | 5.2±0.51$^{bc}$ | 2.9±0.53$^{ab}$ | 1.9±0.42$^{a}$ | 1.2±0.16$^{a}$ | 29.2*** | KW |
| VPD | 2.5±0.08$^{ab}$ | 2.4±0.1$^{a}$ | 2.8±0.12$^{ab}$ | 2.8±0.16$^{ab}$ | 2.4±0.09$^{ab}$ | 3.0±0.18$^{b}$ | 4.4** | ANOVA |

| | Source | | | | | |
|---|---|---|---|---|---|---|
| | Bulk | NP100 | NP50 | NW | value | test |
| SPAD | 38.3±1.6 | 39.7±1.2 | 37.1±1.6 | 40.0±1.9 | 1.3 | KW |
| Fv/Fm | 0.75±0.008$^{a}$ | 0.77±0.006$^{ab}$ | 0.75±0.008$^{a}$ | 0.78±0.008$^{b}$ | 8.4* | KW |
| ΦPSII | 0.13±0.014$^{a}$ | 0.20±0.018$^{c}$ | 0.15±0.021$^{ab}$ | 0.19±0.019$^{bc}$ | 5.5** | ANOVA |
| qP | 0.35±0.027$^{a}$ | 0.47±0.036$^{b}$ | 0.36±0.044$^{ab}$ | 0.44±0.034$^{ab}$ | 10.6* | KW |
| ETR | 68288±7376$^{a}$ | 102297±9244$^{c}$ | 78128±10905$^{ab}$ | 98936±9789$^{bc}$ | 5.5** | ANOVA |

| | | | | | | |
|---|---|---|---|---|---|---|
| qN | 225±122 | 613±193 | 457±182 | 240±163 | 3.9 | KW |
| NPQ | 2392±586 | 1000±405 | 1714±460 | 1590±427 | 4.1 | KW |
| Fv'/Fm' | 2.3±0.06 | 2.3±0.14 | 2.2±0.11 | 2.3±0.17 | 2.9 | KW |
| $\Phi CO_2$ | 0.010±0.002$^{a}$ | 0.015±0.002$^{b}$ | 0.013±0.002$^{ab}$ | 0.015±0.002$^{b}$ | 8.0* | KW |
| A | 9.4±1.7 | 14.6±1.6 | 12.1±1.9 | 14.2±1.9 | 7.8 | KW |
| $g_s$ | 0.12±0.023 | 0.19±0.025 | 0.14±0.024 | 0.19±0.034 | 6.4 | ANOVA |
| E | 2.9±0.42 | 4.3±0.52 | 3.6±0.59 | 4.6±0.62 | 5.8 | KW |
| VPD | 2.7±0.11 | 2.4±0.1 | 2.7±0.08 | 2.7±0.15 | 1.7 | ANOVA |

Plants were grown in four different ZnO sources: micron-size (Bulk), nanoparticles < 100 nm (NP100), nanoparticles < 50 nm (NP50), and nanowires of 50 nm diameter (NW). Each source was provided at six concentrations (0, 0.1, 1, 10, 100, and 1000 mg $l^{-1}$) except NW (only up to 10 mg $l^{-1}$). Data represent means ±SE, where n = 4 and df= 5 for the ZnO concentration factor and df = 3 for the source factor. Variables SPAD, Fv/Fm, ΦPSII, qP, qN, NPQ, Fv'/Fm', and $\Phi CO_2$ are dimensionless. Electron transmission rate (ETR) is expressed in mol $m^{-2}$ $s^{-1}$, net photosynthetic rate (A) in µmol $CO_2$ $m^{-2}$ $s^{-1}$, stomatal conductance (gs) in mol $H_2O$ $m^{-2}$ $s^{-1}$, transpiration (E) in mmol $H_2O$ $m^{-2}$ $s^{-1}$, and vapour pressure deficit of the leaf (VPD) in kPa. Different letters indicate statistically significant groups according to either paired-t-tests with Bonferroni correction (ANOVA) or Dunn's test with Benjamini-Hochberg correction (Kruskal-Wallis, KW). The F-value (ANOVA) or Chi-square value (KW) is indicated as significant at *P* <0.05 (*), *P* <0.01 (**), or *P* <0.001(***).

Supplementary Table 10. Element content in plant tissues.

| | [ZnO] (mg $l^{-1}$) | | | | | | | | Source | | | | | |
|---|---|---|---|---|---|---|---|---|---|---|---|---|---|---|
| | 0 | 0.1 | 1 | 10 | 100 | 1000 | value | test | Bulk | NP100 | NP50 | NW | value | test |
| Root | | | | | | | | | | | | | | |
| [Ca] | 9.8±1.0 | 10.0±0.7 | 10.4±0.6 | 9.1±0.9 | 8.2±1.4 | 8.4±0.9 | 8 | KW | 9.8±0.8 | 8.4±0.7 | 10.1±0.6 | 9.3±0.7 | 3 | KW |
| [Cu] | 22.1±2.3 | 23.6±2.2 | 25.6±1.3 | 25.6±1.6 | 21.2±1.7 | 20.2±1.3 | 9 | KW | 25.8±1.7 | 21.2±1.3 | 21.9±1.3 | 24.5±1.5 | 6 | KW |
| [K] | 17.5±1.1$^{c}$ | 16.6±1.1$^{bc}$ | 15.2±0.8$^{bc}$ | 15.2±1.2$^{bc}$ | 12.3±0.7$^{ab}$ | 10.3±0.7$^{a}$ | 7*** | AV | 14.8±1.0 | 15.5±0.9 | 13.7±0.9 | 15.3±0.9 | 1 | AV |
| [Fe] | 354±28$^{a}$ | 395±34$^{ab}$ | 526±65$^{abc}$ | 583±90$^{bc}$ | 395±32$^{abc}$ | 705±134$^{c}$ | 17** | KW | 598±89 | 464±62 | 460±36 | 423±31 | 2 | KW |
| [Mg] | 1.1±0.1$^{ab}$ | 1.1±0.05$^{ab}$ | 1.3±0.1$^{b}$ | 1.3±0.1$^{ab}$ | 1.1±0.1$^{ab}$ | 1.1±0.1$^{a}$ | 14* | KW | 1.3±0.1 | 1.1±0.1 | 1.2±0.1 | 1.3±0.1 | 5 | KW |
| [Mn] | 30.3±3.4$^{a}$ | 32.4±3.8$^{a}$ | 41.1±4.8$^{ab}$ | 43.8±5.9$^{ab}$ | 44.3±8.2$^{ab}$ | 54.1±4.9$^{b}$ | 15* | KW | 40.5±3.7 | 33.8±3.5 | 49.9±5.4 | 34.7±2.9 | 6 | KW |
| [P] | 6.5±0.3$^{b}$ | 8.2±0.3$^{c}$ | 8.0±0.2$^{c}$ | 7.0±0.3$^{bc}$ | 6.2±0.6$^{ab}$ | 4.9±0.3$^{a}$ | 15*** | AV | 6.9±0.3 | 6.4±0.4 | 7.1±0.3 | 7.3±0.3 | 2 | AV |
| [S] | 3.3±0.1 | 2.9±0.1 | 3.1±0.1 | 3.0±0.1 | 2.9±0.1 | 2.8±0.2 | 10 | KW | 3.1±0.1 | 3.0±0.1 | 2.9±0.1 | 3.0±0.1 | 1 | KW |
| Shoot | | | | | | | | | | | | | | |
| [Ca] | 6.5±0.9 | 6.5±0.7 | 6.0±0.4 | 7.1±0.5 | 7.9±0.7 | 7.1±0.7 | 8 | KW | 7.4±0.7 | 6.4±0.4 | 6.5±0.4 | 6.9±0.7 | 1 | KW |
| [Cu] | 12.5±1.0$^{b}$ | 17.6±1.5$^{c}$ | 15.8±0.7$^{c}$ | 13±0.9$^{bc}$ | 10.3±0.5$^{ab}$ | 8.8±0.7$^{a}$ | 42*** | KW | 13.9±1.0 | 13.1±1.3 | 11.9±0.6 | 15.1±1.0 | 7 | KW |
| [K] | 32.5±1.2$^{a}$ | 32.4±0.9$^{a}$ | 32.6±0.9$^{a}$ | 36.0±0.9$^{b}$ | 37.2±1.8$^{b}$ | 32.4±1.5$^{a}$ | 3* | AV | 33.5±0.9 | 34.7±1.1 | 33.8±0.9 | 32.2±1.2 | 1 | AV |
| [Fe] | 98.7±12.9 | 89.5±7.1 | 91.4±5.0 | 102.1±12.1 | 68.4±5.2 | 92.1±9.7 | 9 | KW | 96.7±9.7 | 83.1±6.3 | 87.1±5.6 | 102.1±9.7 | 3 | KW |
| [Mg] | 1.7±0.1 | 1.7±0.1 | 1.6±0.1 | 1.6±0.1 | 1.8±0.1 | 1.6±0.1 | 1 | KW | 1.7±0.1 | 1.7±0.1 | 1.6±0.1 | 1.7±0.1 | 0 | KW |
| [Mn] | 101.2±12.7$^{b}$ | 112.9±16.2$^{b}$ | 109.9±11.3$^{b}$ | 54.7±3.2$^{a}$ | 35.2±3.3$^{a}$ | 34.3±3.3$^{a}$ | 50*** | KW | 79.1±10.3$^{ab}$ | 61.5±8.3$^{a}$ | 69.7±9.0$^{a}$ | 117.8±16.3$^{b}$ | 12** | KW |
| [P] | 3.2±0.3$^{a}$ | 3.0±0.1$^{a}$ | 3.3±0.1$^{ab}$ | 4.3±0.3$^{c}$ | 4.3±0.2$^{c}$ | 3.7±0.2$^{bc}$ | 37*** | KW | 3.8±0.2 | 3.5±0.2 | 3.7±0.2 | 3.4±0.2 | 2 | KW |
| [S] | 7.6±0.4$^{a}$ | 8.3±0.4$^{ab}$ | 8.4±0.3$^{ab}$ | 9.7±0.5$^{b}$ | 10.1±0.7$^{b}$ | 7.3±0.5$^{a}$ | 27*** | KW | 8.8±0.4 | 8.5±0.4 | 8.2±0.4 | 8.5±0.4 | 1 | KW |

Concentrations are provided in mg $g^{-1}$ except for Cu, Fe, and Mn in roots (in µg $g^{-1}$). Plants were grown in four different ZnO sources: micron-size (Bulk), nanoparticles < 100 nm (NP100), nanoparticles < 50 nm (NP50), and nanowires of 50 nm diameter (NW). Each source was provided at six concentrations (0, 0.1, 1, 10, 100, and 1000 mg $l^{-1}$) except NW (only up to 10 mg $l^{-1}$). Data represent means ±SE, where n = 4 and df= 5 for the ZnO concentration factor and df = 3 for the source factor. Different letters indicate statistically significant groups according to either paired-t-tests with Bonferroni correction (ANOVA) or Dunn's test with Benjamini-Hochberg correction (Kruskal-Wallis, KW). The F-value (ANOVA) or Chi-square value (KW) is indicated as significant at $P$ < 0.05 (*), $P$ <0.01 (**), or $P$ <0.001(***).

Supplementary Fig. 1. Distribution of Al across the different pools. Plants were treated with four different ZnO sources: A) micron-size (Bulk), B) NP < 100 nm (NP100), C) NP < 50 nm (NP50), and D) nanowires of 50 nm diameter (NW). Data represent means, where n = 4, expressed as Al % relative to the total Al incorporated to the system from the nutritive solution and ZnO treatments.

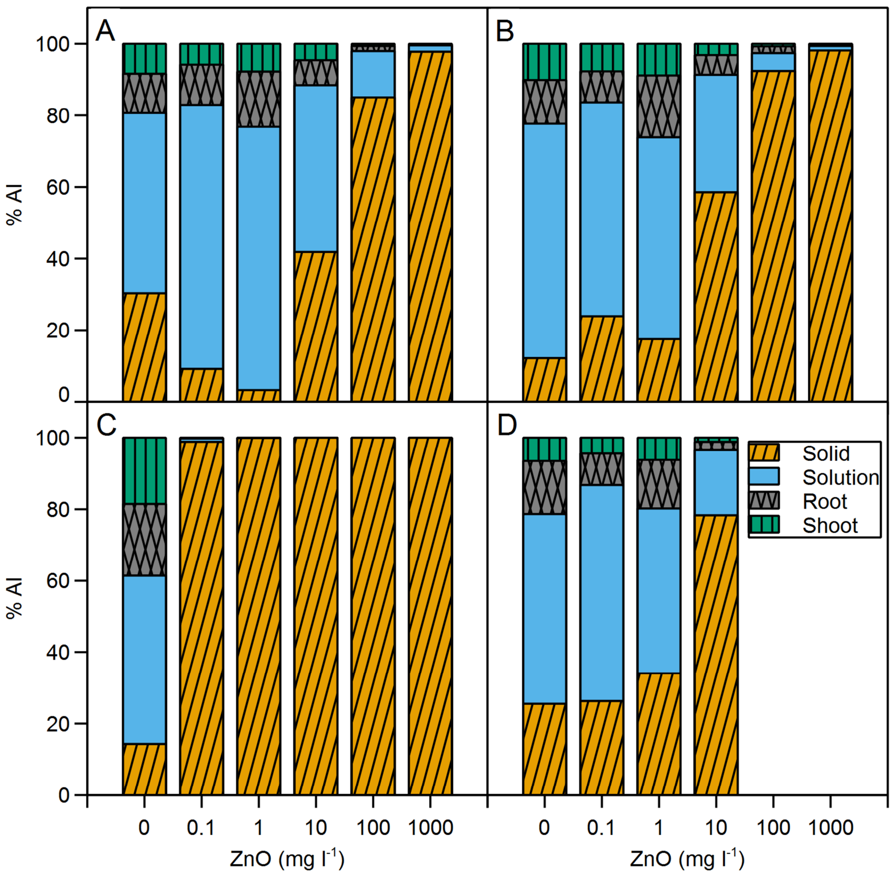

Supplementary Fig. 2. Light microscopy images of cross 1 µm sections of the root. A) Control; B) Bulk, 1000 mg $l^{-1}$ ZnO; C) NP100, 1000 mg $l^{-1}$ ZnO; D) NP50, 1000 mg $l^{-1}$ ZnO; E) NW, 10 mg $l^{-1}$ ZnO.

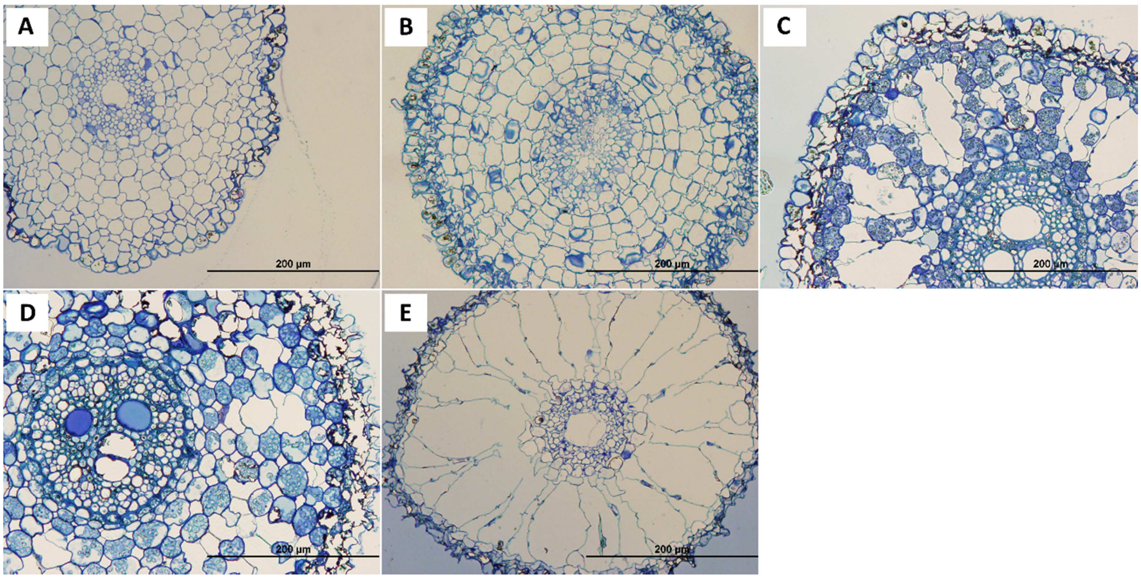

Supplementary Fig. 3. Transmission electron microscopy images of the root epidermis. A) Control; B) Bulk, 1000 mg $l^{-1}$ ZnO; C) NP100, 1000 mg $l^{-1}$ ZnO; D) NP50, 1000 mg $l^{-1}$ ZnO.

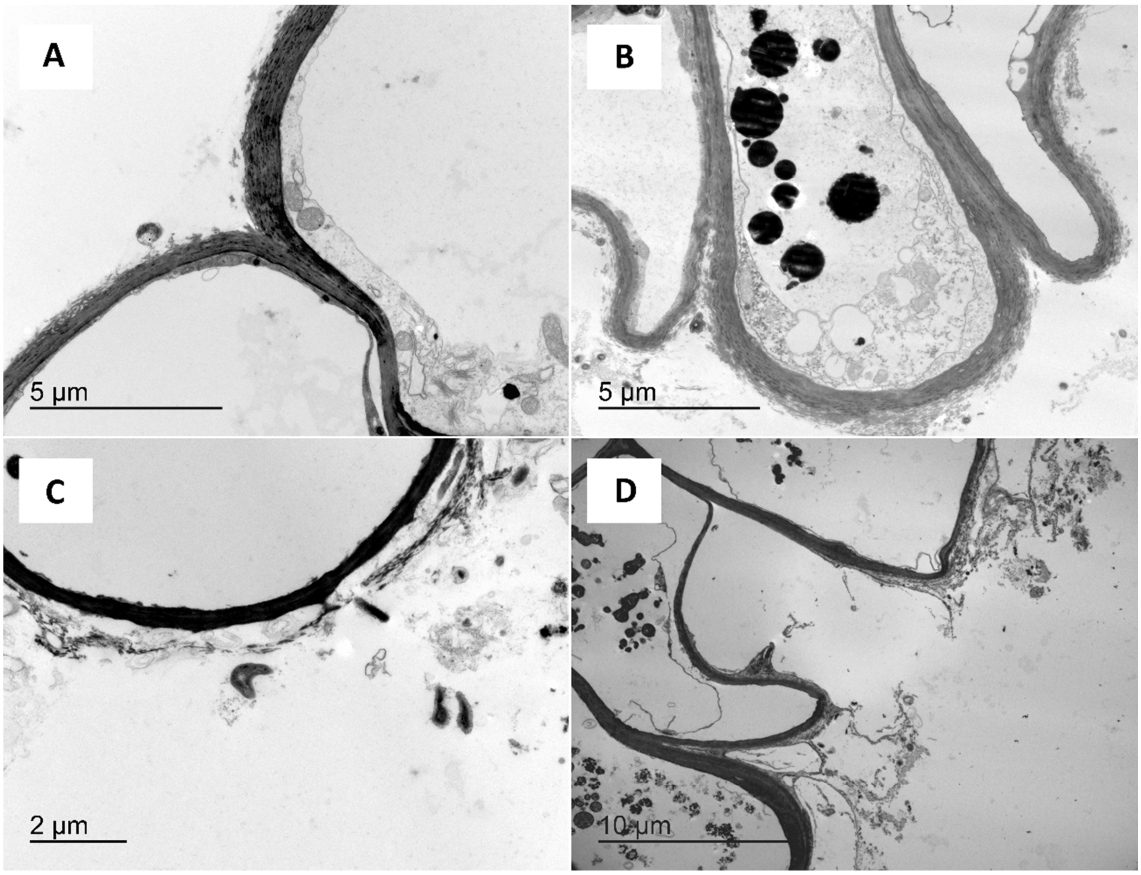

Supplementary Fig. 4. Transmission electron microscopy images of the root cortex. A) Control; B) Bulk, 1000 mg $l^{-1}$ ZnO; C) NP100, 1000 mg $l^{-1}$ ZnO; D) NP50, 1000 mg $l^{-1}$ ZnO.

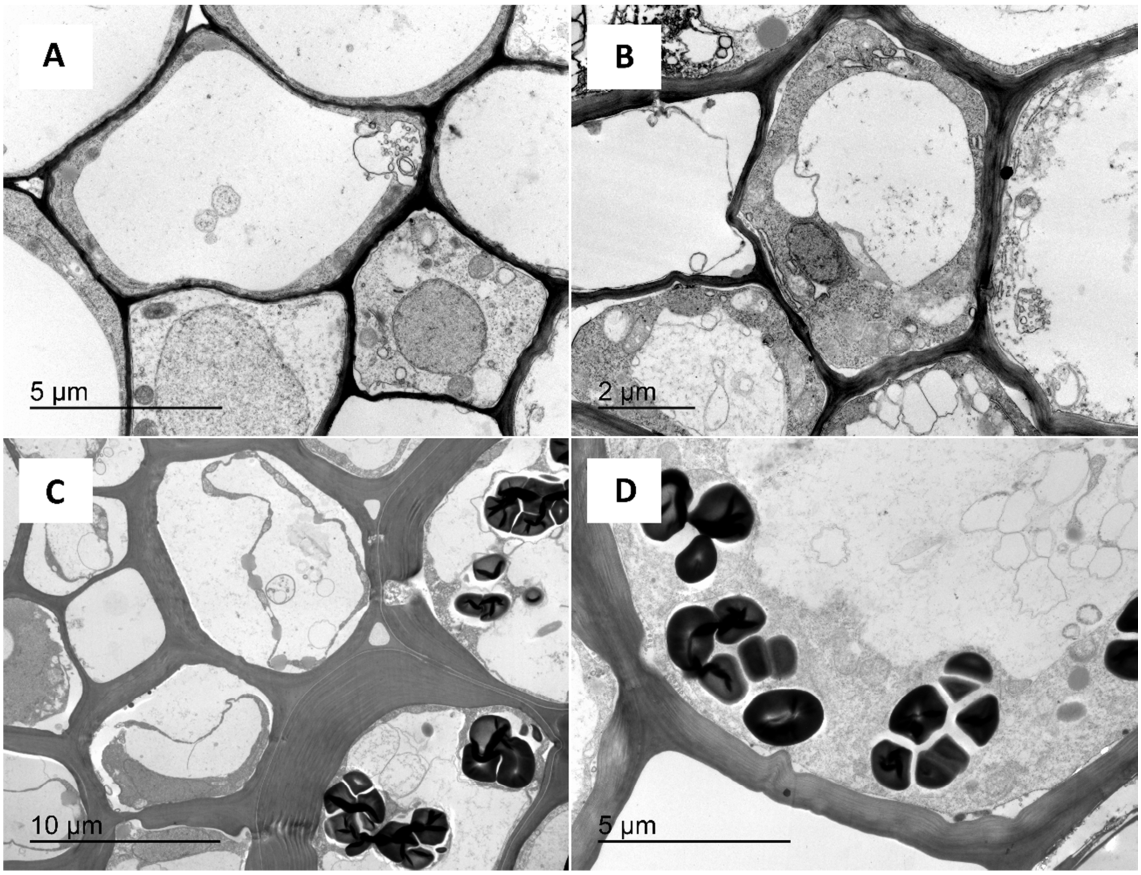
